# Mechanics of human embryo compaction

**DOI:** 10.1101/2022.01.09.475429

**Authors:** Julie Firmin, Nicolas Ecker, Diane Rivet Danon, Virginie Barraud Lange, Hervé Turlier, Catherine Patrat, Jean-Léon Maître

**Affiliations:** Institut Curie, Université PSL, CNRS UMR3215, INSERM U934, 75005 Paris, France; Université de Paris, Paris, France; Center for Interdisciplinary Research in Biology, Collège de France, CNRS, INSERM, Université PSL, FHU Prema, Paris, France; Service de Biologie de la Reproduction - CECOS, Paris Centre Hospital, APHP centre, FHU Prema, 75014; Institut Cochin, Université de Paris, CNRS UMR1016, 75014 Paris, France

## Abstract

The shaping of the human embryo begins with compaction, during which cells come into close contact and form a tighter structure^1,2^. Assisted reproductive technology (ART) studies suggest that human embryos fail compaction primarily because of defective adhesion^3,4^. Based on our current understanding of animal morphogenesis^5,6^, other morphogenetic engines, such as cell contractility, could be involved in shaping the human embryo. However, the molecular, cellular and physical mechanisms driving human embryo morphogenesis remain uncharacterized. Using micropipette aspiration on human embryos donated to research, we have mapped cell surface tensions during compaction. This reveals a 4-fold increase of tension at the cell-medium interface while cell-cell contacts keep a steady tension. Comparison between human and mouse reveals qualitatively similar but quantitively different mechanical strategies, with human embryos being mechanically least efficient. Inhibition of cell contractility and cell-cell adhesion in human embryos reveal that only contractility controls the surface tension responsible for compaction. Interestingly, if both cellular processes are required for compaction, they exhibit distinct mechanical signatures when faulty. Analyzing the mechanical signature of naturally failing embryos, we find evidence that non-compacting embryos or partially compacting embryos with excluded cells have defective contractility. Together, our study reveals that an evolutionarily conserved increase in cell contractility is required to generate the forces driving the first morphogenetic movement shaping the human body.

---

The mechanical characterization of model organisms, including mammals, has immensely advanced our understanding of animal morphogenesis^5,6^. For ethical and technical reasons, human embryos are mostly inaccessible to experimentation. Therefore, our appreciation of how the human body shapes itself during embryonic development rarely comes from studies on human embryos themselves, but instead relies mostly on the extrapolation from findings in other species and, more recently, from engineered human embryo models^7,8^. For example, we still do not know whether contractility of the actomyosin cortex, a major morphogenetic engine during animal development^5,6,9^, plays a similarly important role during human morphogenesis^1^. Opportunely, preimplantation development constitutes a unique setting to carry out experimentations on live embryos and can provide both validation and breakthrough in our understanding of human embryonic development^10–13^.

Human morphogenesis begins with compaction on the 4^th^ day after fertilization, when the embryo is composed of 8 to 16 cells^1–3,14^. After *in vitro* fertilization (IVF) during ART, embryos failing to compact entirely or with a delayed compaction show lower implantation rate upon transfer^15–17^, illustrating the importance of this process for further development. Also, human embryos can compact partially, with individual cells being excluded from the compacted mass^3,4^. However, the mechanisms leading to compaction failure in human embryos are unknown.

During compaction, cells maximize their cellcell contact area and minimize their surface exposed to the outside medium^18^. This is akin to the adhesion of soap bubbles resulting from the balance of tensions at their interfaces. Following this analogy and as described previously^19,20^, we consider the surface tensions *γ*_cc_ and *γ*_cm_ at cell-cell contacts and at cell-medium interfaces respectively, whose ratio determines the shape of contacts between cells. Precisely, compaction of human embryos results from reducing a compaction parameter α = cos ( θ_e_ / 2) = *γ*_cc_ / 2 *γ*_cm_, where θ_e_ is the external contact angle between cells (Supplementary Information, Fig 1a). Using time-lapse microscopy and micropipette aspiration, we have determined the contact angles and surface tensions of human embryos (Fig 1a-d, Movie 1). Synchronizing embryos to the last observed 3^rd^ cleavage, we measured the growth of external contact angles θ_e_ from 81 ± 5 to 158 ± 4° in ~35 h (mean ± SEM of 149 measurements on 10 embryos, Student’s t test p < 10-^11^, Fig 1c, Table 1). Concomitantly, surface tensions *γ*_cm_ increase from 615 ± 39 to 2347 ± 84 pN/*μ*m (mean ± SEM of 147 measurements on 10 embryos, Student’s t test p < 10^-5^) while surface tensions at cell-cell contacts *γ*_cc_/2 remained steady at ~600 pN/*μ*m (Fig 1d, Table 1). Independently of embryo synchronization, calculating the correlation between contact angles and surface tensions yields 0.625 for *γ*_cm_ and −0.134 for *γ*_cc_ over the entire duration of the experiments (433 and 196 measurements on 14 embryos and Pearson correlation p values < 10^-47^ and > 10^-2^ respectively, Fig 1e-f). Therefore, the mechanical changes driving compaction are located at the cell-medium interface rather than at cell-cell contacts (Fig 1).

**Figure 1:**
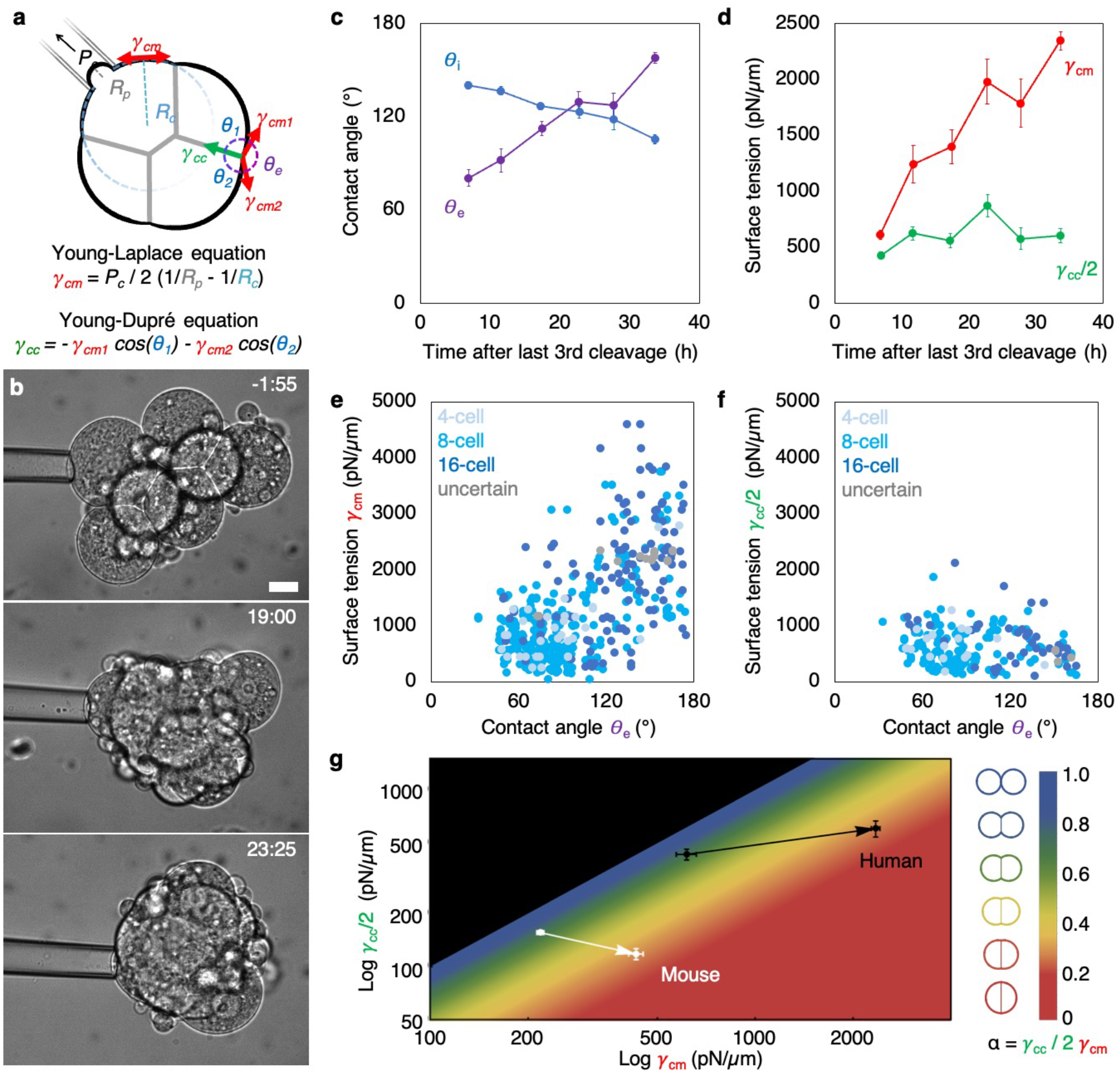
Spatiotemporal map of tensions during human embryo compaction. (a) Diagram of the tension mapping method. Using a micropipette of radius *R_p_*, a pressure *P_c_* is applied to the surface of blastomeres of curvature 1/*R_c_*. The surface tension *γ*_cm_ is calculated using the Young-Laplace equation. From *γ*_cm_, the external and internal contact angles (θ_e_ and, θ_1_ and θ_2_, respectively) of adjacent cells, we use the Young-Dupré equation to calculate the interfacial tension *γ*_cc_. (b) Representative images of human embryos during micropipette aspiration. Time relative to last observed 3^rd^ cleavage division as hh:mm. Scale bar, 20 *μ*m. Movie 1. (c-d) Time course of internal and external contact angles θ_1_ and θ_e_ respectively (c) and surface tensions *γ*_cm_ and *γ*_cc_/2 (d). Mean ± SEM over 5 h of 147 blastomeres and 103 contacts from 10 embryos synchronized to the time of last observed 3^rd^ cleavage division. Internal contact angles θ_1_ and θ_2_ are averaged to calculate θ_1_. Statistics in Table 1. (e-f) Surface tension *γ*_cm_ and *γ*_cc_/2 as a function of contact angles θ_e_ measured on 257 contacting blastomeres from 14 embryos. Pearson correlation values R = 0.625 for *γ*_cm_ (p < 10^-47^) and R = −0.134 for *γ*_cc_ (p > 10^-2^). Cleavage stages are indicated with 4-, 8- and 16-cell stage blastomeres in light, medium and dark blue respectively. Grey dots show blastomeres that cannot be staged with certainty. (g) Phase diagram showing the state of compaction as a function of *γ*_cm_ and *γ*_cc_/2 in log-log scale. Mean ± SEM of data from human embryos are shown as a black arrow starting at 5-10 and ending at 30-35 h after the last 3^rd^ cleavage. Mean ± SEM of data from mouse embryos adapted from (Maître et al, 2015)^20^ are shown as a white arrow starting at 0-2 and ending at 8-10 h after the 3^rd^ cleavage. The compaction parameter α = *γ*_cc_ / 2 *γ*_cm_ is colour-coded on the right, with diagrams of the corresponding cell doublet shapes.

As cleavage divisions of blastomeres within individual embryos can occur hours apart from one another, compaction takes place with neighboring cells at different cleavage stages (Fig 1e-f, Movie 1). Tracking cleavage stages, surface tensions and contact angles revealed that the mechanical changes associated with compaction can begin during the 8-cell stage and proceed as blastomeres undergo their 4^th^ cleavage (Fig 1e-f). This is delayed compared to the mouse, in which the mechanical changes driving a compaction of similar magnitude occur during the 8-cell stage^20^. Further comparison between mouse and human reveals that the increase in surface tension *γ*_cm_ during compaction is qualitatively conserved (Fig 1g). However, while mouse embryos double their surface tension *γ*_cm_, human embryos increase it 4-fold to drive contact angle changes of identical magnitude (Fig 1g, Supplementary Note). In addition to increasing *γ*_cm_, compaction in mouse embryos also relies on decreasing *γ*_cc_ (Fig 1g). We had previously calculated that, in the mouse changes in tension *γ*_cm_ contributed to three-quarter of compaction and changes in tension *γ*_cc_ to one-quarter^20^. This is not the case in human embryos, which do not relax their cellcell contacts and rely exclusively on the increase in tension *γ*_cm_ at the cell-medium interface (Fig 1g). Therefore, mouse and human embryos share qualitatively conserved mechanisms but quantitatively different strategies to achieve the same morphogenesis. Interestingly, considering the minimal changes in surface tension required to compact, the strategy adopted by human embryos is less efficient than the one of the mouse (Fig 1g, Supplementary Note). Indeed, with growing external contact angles, any further increase in tension *γ*_cm_ becomes less and less effective due to cells pulling more and more perpendicularly to the plane of cell-cell contacts^18,20^. Therefore, compared to the mouse, human embryos must generate considerable surface stresses with unknown implications for human embryo development (Fig 1g).

In mouse embryos, the increase in tension *γ*_cm_ is mediated by the contractility of the actomyosin cortex and the reduction in tension *γ*_cc_ results in part from the downregulation of contractility, which requires signals from cadherin adhesion molecules^20,21^. To investigate the molecular and cellular regulation of the mechanics of human embryo compaction, we analyzed the distribution of filamentous actin (F-actin), non-muscle myosin 2B (MYH10) and E-cadherin (CDH1, Fig 2a). We found that, as contact angles grow, MYH10 becomes enriched at the cell-medium interface as compared to cell-cell contacts while F-actin and CDH1 levels remain stable (Fig 2a-b). This is compatible with increased contractility at the cell-medium interface, underlying raising tension *γ*_cm_, and steady cell-cell contacts, associated with stable *γ*_cc_ (Fig 1d).

**Figure 2:**
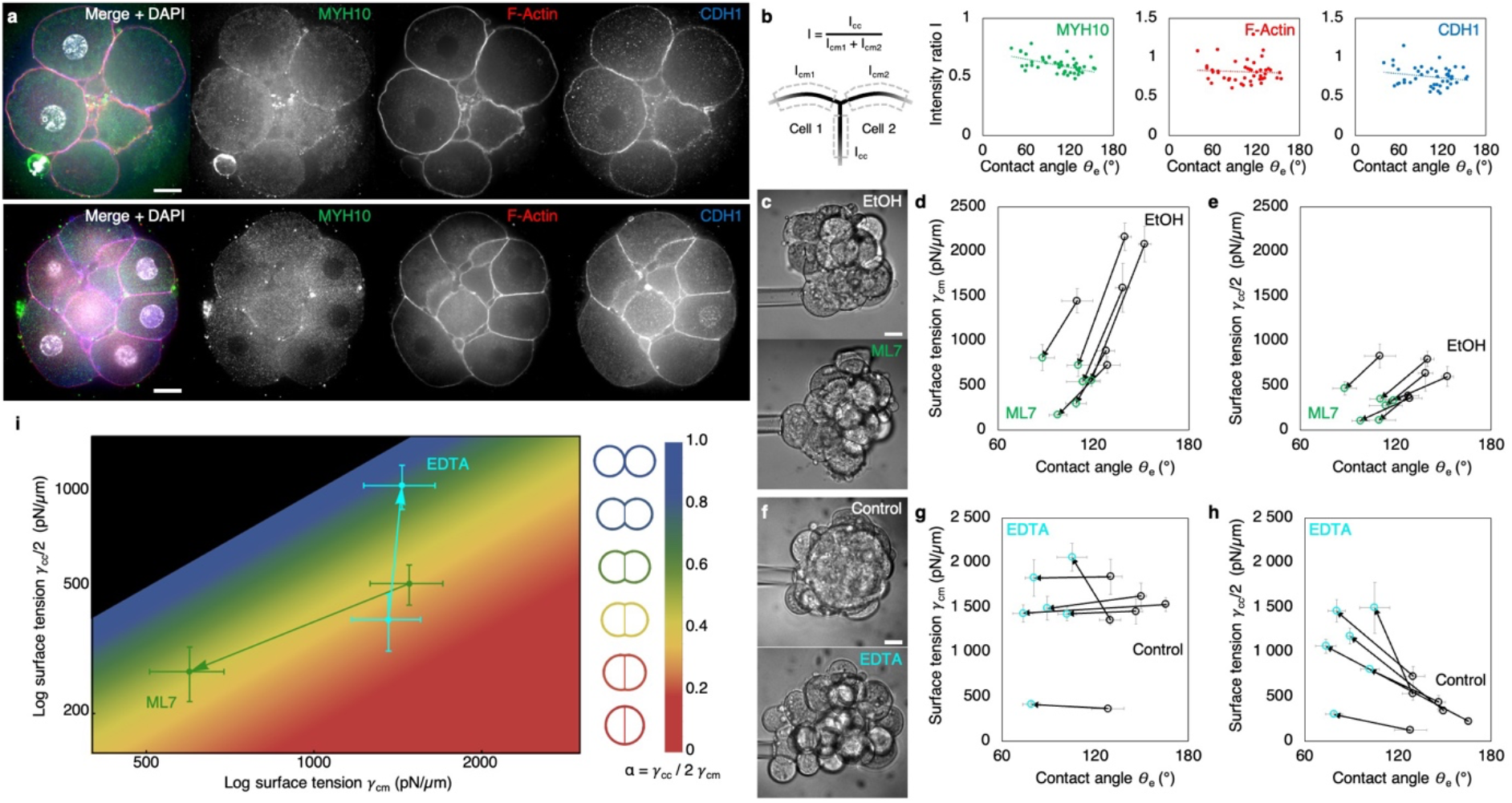
Role of cell contractility and cell adhesion in regulating tensions during human embryo compaction. (a) Representative images of immunostaining of human embryos before (top) and after (bottom) compaction with MYH10 in green, F-actin in red and CDH1 in blue. Scale bar, 20 *μ*m. (b) Intensity ratio between the cell-cell and cell-medium interfaces for MYH10 in green, F-actin in red and CDH1 in blue as a function of the external contact angle θ_e_. Pearson correlations R = −0.535 for MYH10 (p < 10^-3^), −0.069 for F-actin (p > 10^-1^) and −0.203 for CDH1 (p > 10^-1^). 45 cells from 10 embryos. (c) Representative images of embryos placed in medium containing 1:2000 EtOH (top) and 10 μM ML7 (bottom) during micropipette aspiration. Scale bar, 20 *μ*m. (d-e) Surface tension *γ*_cm_ (d) or *γ*_cc_ (e) as a function of external contact angles θ_e_ measured on 115 blastomeres from 6 embryos placed in EtOH (black) and ML7 (green) media. Comparisons between EtOH and ML7 media using pairwise Student’s t test give p < 10^-4^ for θ_e_, p < 10^-2^ for *γ*_cm_ and p < 10^-3^ for *γ*_cc_/2. Statistics in Table 2. (f) Representative images of embryos placed in control (top) and EDTA (bottom) media during micropipette aspiration. Scale bar, 20 *μ*m. (g-h) Surface tension *γ*_cm_ (g) or *γ*_cc_/2 (h) as a function of external contact angles θ_e_ measured on 98 blastomeres from 6 embryos placed in normal (black) and EDTA (blue) media. Comparisons between control and EDTA media using pairwise Student’s t test give p < 10^-2^ for θ_e_, p > 10^-1^ for *γ*_cm_ and p < 10^-3^ for *γ*_cc_/2. Statistics in Table 2. (i) Phase diagram showing the state of compaction as a function of *γ*_cm_ and *γ*_cc_/2 in log-log scale. Mean ± SEM of data from embryos transferred from EtOH to ML7 media (green, 60, 55 cells from 6 embryos) or from control to EDTA media (blue, 49, 49 cells from 6 embryos). The compaction parameter α = *γ*_cc_ / 2 *γ*_cm_ is colour-coded on the right, with diagrams of the corresponding cell doublet shapes.

To test whether contractility is responsible for generating the tensions driving human embryo compaction, we used ML7, an inhibitor of the myosin light chain kinase, on compacted embryos. ML7 caused embryos to decompact with contact angles dropping by 27 ± 2° within minutes (mean ± SEM from 6 embryos, pairwise Student’s t test p < 10^-4^, Fig 2c-e, Table 2). Concomitantly, we measured a 3-fold decrease in tension *γ*_cm_ between embryos in control and ML7-containing media (pairwise Student’s t test p < 10^-2^, Fig 2d, Table 2). Importantly, placing embryos back in regular medium allowed embryos to compact again (6/6 embryos) and to form a blastocyst (5/6 embryos). This indicates that contractility is required for generating *γ*_cm_, as in mouse embryos^20^. Therefore, contractility is an evolutionarily conserved engine generating the tension *γ*_cm_ driving compaction of both human and mouse embryos. Upon ML7 treatment, the tension at cell-cell contacts *γ*_cc_ also decreases by half (pairwise Student’s t test p < 10^-3^, Fig 2e, Table 2), indicating high levels of contractility acting at cell-cell contacts of human embryos. This is different from what has been reported in mouse embryos, which display minimal contractility at their cell-cell contacts^20^. Since reducing *γ*_cc_ promotes compaction, high levels of contractility at cell-cell contacts also explain why global inhibition of contractility shows milder effects on compaction in human embryos as compared to mouse ones. Furthermore, high levels of contractility at cellcell contacts could explain why human embryos increase their tension *γ*_cm_ twice as much as mouse ones in order to compact.

Despite lacking obvious molecular reorganization during compaction (Fig 2a-b), cadherin-based cell-cell adhesion remains the prime suspect for compaction defects observed in ART^3,4^. Therefore, we decided to investigate the influence of cadherin-based adhesion onto surface tension in human embryos. Since cadherin adhesion molecules require Ca^2+^ to function^22^, we placed compacted embryos in medium without Ca^2+^ and supplemented with EDTA (Fig 2f). As previously observed^23^, EDTA medium led to rapid decompaction of human embryos with contact angles dropping by 53 ± 8° (mean ± SEM from 6 embryos, pairwise Student’s t test p < 10^-2^, Fig 2g-h, Table 2). As for contractility inhibition, placing embryos back into regular medium allowed embryos to compact again (6/6 embryos) and to form a blastocyst (4/6 embryos). Surface tension measurements revealed that tension *γ*_cm_ was not affected by EDTA medium while tension *γ*_cc_ increased 2-fold (pairwise Student’s t test p > 10^-1^ and p < 10^-2^ respectively, Fig 2g-h, Table 2). Therefore, as observed in mouse embryos^20^, the tension *γ*_cm_ driving compaction is generated independently from cell adhesion, in a cell-autonomous manner. Embryos lacking adhesion decompact because of cells’ inability to transmit tensions to their neighbors. The increase in tension *γ*_cc_ may result from loss of adhesion energy from cadherin adhesion molecules binding^22^ or, as observed in the mouse, from upregulation of contractility at cellcell contacts^20,24^.

Together, we find that, while both cell contractility and cell-cell adhesion are required for compaction of human embryos (Fig 2c-h), only contractility reorganizes during compaction and generates the tension *γ*_cm_ that drives compaction (Fig 2a-b). Therefore, human embryo compaction relies on contractility increasing surface tension specifically at the cell-medium interface, which constitutes molecular, cellular and mechanical mechanisms that are qualitatively conserved with the mouse embryo.

Interestingly, loss of contractility and adhesion results in distinct mechanical signatures (Fig 2i): low *γ*_cm_ and *γ*_cc_ with ML7 medium in contrast to high *γ*_cm_ and *γ*_cc_ with EDTA medium. Therefore, mechanical signatures could be used to distinguish which cellular process fails when compaction is defective and potentially help diagnostics in IVF clinics^1,25^.

To determine the mechanical origin of compaction defects, we measured the tensions of embryos spontaneously failing compaction. We considered compaction failed when no contact angle would grow above 131°, as determined statistically from 7 compacting and 7 non-compacting embryos^26^. For these embryos, 30 h after the 3^rd^ cleavage and despite cells undergoing their 4^th^ cleavage similarly to compacting embryos (Extended Data Fig 1), mean contact angles kept steady below ~80° (Fig 3a-b, Table 3). Meanwhile, both tensions *γ*_cm_ and *γ*_cc_ remained low (Fig 3b- d, Table 3). This corresponds to the mechanical signature of defective contractility (Fig 2i), indicating that all of the 7 spontaneously failing embryos we have measured were unsuccessful in growing their contractility. Therefore, defective contractility could be a common cause of compaction failure in human embryos. Importantly, if both contractility and adhesion were defective, we would expect low tensions, which would be indistinguishable from faulty contractility alone. Therefore, in addition to contractility, adhesion may also be deficient in the embryos we have measured.

**Figure 3:**
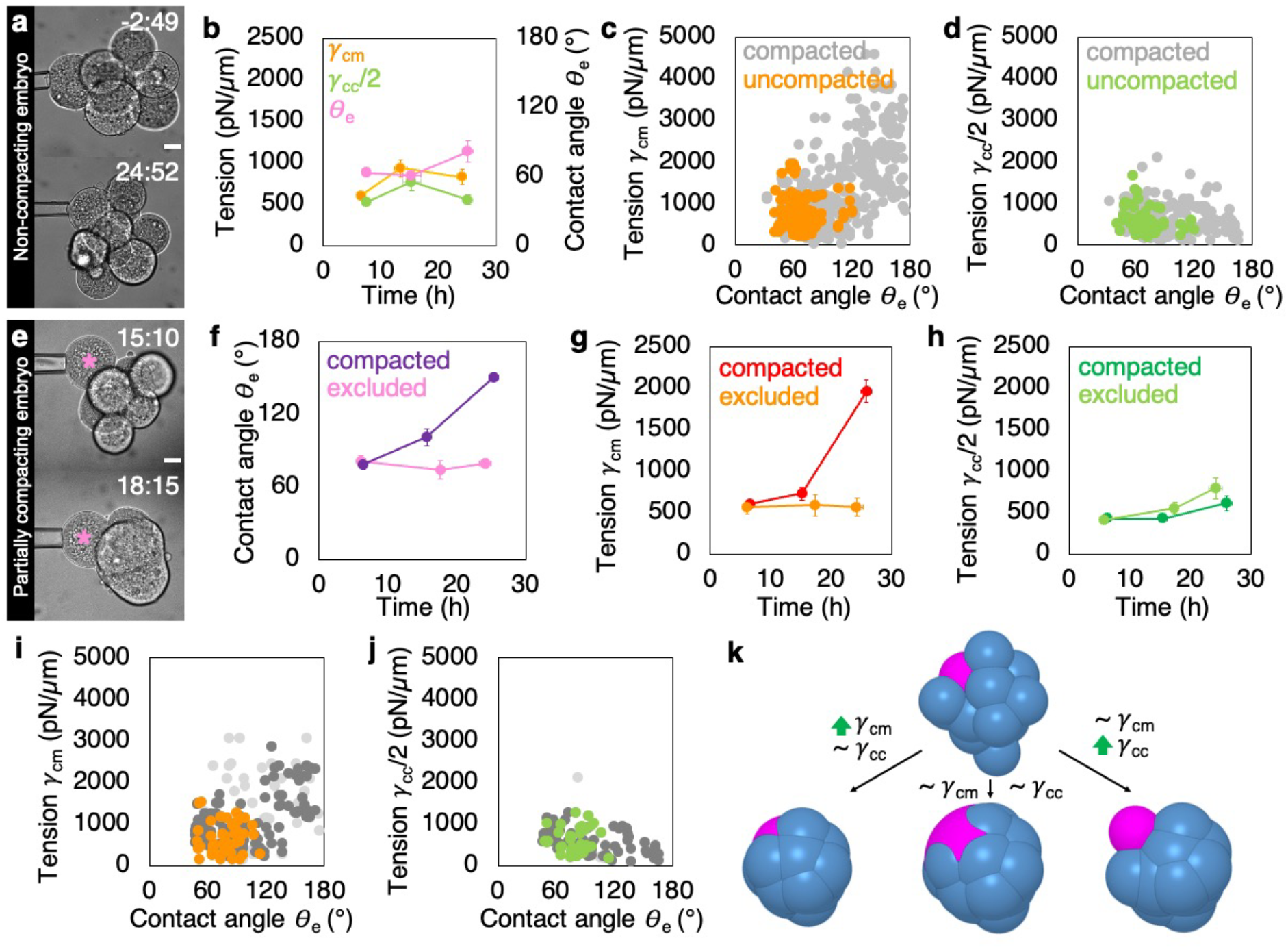
Mechanical signature of human embryos failing compaction. (a) Representative images of embryos failing compaction with no contact reaching 131 ° for over 30 h after the 3^rd^ cleavage. Time relative to last observed 3^rd^ cleavage division as hh:mm. Scale bar, 20 *μ*m. Movie 2. (b) Time course of contact angles θ_e_ (pink) and surface tensions *γ*_cm_ (orange) and *γ*_cc_/2 (light green). Mean ± SEM over 10 h of 31 blastomeres and 16 contacts from 5 non-compacting embryos synchronized to the time of last observed 3^rd^ cleavage division. Statistics in Table 3. (c-d) Surface tension *γ*_cm_ (c) and *γ*_cc_/2 (d) as a function of contact angles θ_e_ measured on 117 blastomeres from 7 non-compacting embryos. Data from compacting embryos from Fig 1e-f shown in grey. (e) Representative images of a blastomere (labeled with pink asterisk) becoming excluded during compaction with contact angles < 113° within compacting embryos with contact growing above 131° for over 30 h after the 3^rd^ cleavage. Time relative to last observed 3^rd^ cleavage division as hh:mm. Scale bar, 20 *μ*m. Movie 3. (f-h) Time course of external contact angles θ_e_ (f) and surface tensions *γ*_cm_ (g) and *γ*_cc_/2 (h) of compacting (purple, red and dark green) and non-compacting (pink, orange and light green) blastomeres. Mean ± SEM over 10 h of 65 blastomeres and 90 contacts from 5 embryos synchronized to the time of last observed 3^rd^ cleavage division. Statistics in T able 4. (i-j) Surface tension *γ*_cm_ (i) and *γ*_cc_/2 (j) as a function of external contact angles θ_e_ measured on 224 blastomeres from 7 partially compacted embryos. Data from excluded blastomeres in orange (i) and light green (j), compacting blastomeres neighboring excluded cells in light grey and in dark grey for other compacting blastomeres. (k) Simulations of compaction with distinct cell populations: blue blastomeres grow their tension *γ*_cm_ by a factor 3.2 and their tension *γ*_cc_ by 1.4 according to measurements shown in Fig 3g-h; purple blastomeres do the same as blue ones (bottom left), keep *γ*_cm_ steady and grow their tension *γ*_cc_ by 1.4 (bottom middle) or keep *γ*_cm_ steady and grow their tension *γ*_cc_ by 2.4. Movie 4.

Another compaction defect that is commonly observed is partial compaction, where some of the blastomeres do not participate in the compacted mass^1,3,4^. Such excluded cells are thought to be either eliminated from the blastocyst or could participate to extra- embryonic tissues such as the trophectoderm that forms at the surface of the embryo^1,2,27^. Biopsy of excluded cells suggests that those are more likely to be aneuploid, which led to the interesting hypothesis that compaction would serve as a way to eliminate aneuploid cells from embryonic tissues^4^. Furthermore, recent clonal analyses on human placenta from natural pregnancies found that aneuploid clones originate from blastomeres that had segregated into the trophectoderm during preimplantation development^27^. This further supports the idea that human preimplantation embryos eliminate aneuploid cells from embryonic tissues. However, how defective cells would be eliminated from the compacting morula is unknown.

To investigate this mechanism, we measured the tension of embryos showing partial compaction (Fig 3e). We considered cells as excluded when they failed to raise their contact angle θ_e_ above 113° while the rest of the embryo compacted, as determined statistically from 7 embryos containing 1 or more excluded cells^26^. Within the same embryo, compacting blastomeres showed increasing contact angles θ_e_ and tensions *γ*_cm_ while contact tensions *γ*_cc_ would remain steady, as described above (Fig 1d and 3f, Table 4). Meanwhile, non-compacting blastomeres kept their contact angles θ_e_ and tension *γ*_cm_ low, ending up excluded from the compacted morula with minimal attachment (Fig 3f-j, Table 4). Thus, non-compacting blastomeres seem to lack contractility, similarly to embryos failing compaction entirely (Fig 3b-c). Furthermore, the sorting out of non-compacting cells based on differences in tension *γ*_cm_ is reminiscent of the mechanism driving the positioning of low *γ*_cm_ trophectoderm progenitors and high *γ*_cm_ inner cell mass progenitors in the mouse embryo^28^. However, contrary to cells with low tensions *γ*_cm_ that sort out to the surface to become TE cells, excluded cells do not spread at the surface and instead keep minimal attachment (Fig 3e). To understand how cells could become excluded, we simulated different scenarios of surface tension changes using a 3D active foam model of the embryo (Supplementary Note)^28^. Using measured values of *γ*_cm_ and *γ*_cc_ faithfully recapitulated normal compaction *in silico* (Fig 3k, Movie 4). To consider cell exclusion, we maintained *γ*_cm_ at its initial levels in one cell, while changing *γ*_cm_ normally in neighboring cells as measured experimentally (Fig 3g). When both *γ*_cm_ and *γ*_cc_ were maintained low, we did not observe exclusion but noted instead the spreading of the weak cell (Fig 3k, Movie 4). On the other hand, increasing *γ*_cc_ led to cell exclusion (Fig 3k, Movie 4), as observed experimentally. Therefore, exclusion requires *γ*_cc_ to increase (Fig 3k). Indeed, unlike for compacting cells, we measured a 2-fold increase in *γ*_cc_ between excluded cells and their compacting neighbor (from 417 ± 46 to 797 ± 126 pN/*μ*m for *γ*_cc_/2 in 5 embryos, mean ± SEM, Student’s t test p < 0.04, Fig 3h, Table 4). This increase could arise from high contractility at cell-cell contacts from both or from only one of the contacting cells. Since our measurement of *γ*_cm_ suggest that excluded cells have low contractility, high *γ*_cc_ is more likely to originate from the contractility of the neighboring non-excluded cells (Fig 3j). Increased contractility specifically at this interface would constitute an active mechanism by which cells from human embryos would recognize and eliminate unfit cells. Investigating this mechanism further will benefit from the rich literature on cell competition reported in model organisms^29^.

Together, using the mechanical signatures of human preimplantation embryos^25^, we can provide a more accurate explanation for compaction defects that are commonly observed in ART studies^4,16^. Moreover, this first study on the mechanics of human embryonic morphogenesis reveals that normal compaction results from an evolutionarily conserved increase in cell contractility (Fig 1-2). Although qualitatively conserved, the force patterns driving the same morphogenetic movement in mouse and human are quantitively different, with human embryos being less mechanically efficient than mouse ones (Supplementary Note, Fig 1g). Therefore, we uncover that the same morphogenesis does not necessarily rely on identical force patterns, which could be reminiscent of developmental system drift reported for signaling modules^30^. We think this illustrates how studying the evolution of morphogenesis immensely benefits from our ability to measure mechanical properties of embryos in ways that allow comparison, ideally directly with a physical unit^31^. This will be key to discover how physical laws are used by nature to produce the breathtaking diversity of the shapes of life.

## Acknowledgements

We thank the imaging platform of the Genetics and Developmental Biology unit at the Institut Curie (PICT-IBiSA@BDD), member of the French National Research Infrastructure France-BioImaging (ANR-10-INBS-04) for their outstanding support. We thank Nadia Kazdar, Lucie Delaroche and Anne Le Dû and all the members of ART teams from the Clinique La Muette (Paris, France), the Clinique Pierre Chérest (Neuilly sur Seine, France) and the Hopital Cochin (Paris, France) for support with human embryo experiments. We thank all members of the Maître lab, Y. Bellaïche and M.- H. Verlhac for discussion and comments. We acknowledge the support with administrative issues from M.-H. Verlhac throughout this project. We are grateful to the patients who donated their surplus embryos to research.

This project was funded by a Paris Sciences et Lettres (PSL) QLife (ANR-17-CONV-0005) grant to J.-L.M., C.P. and H.T. and the INSERM transversal program Human Development Cell Atlas (HuDeCA) to J.-L.M. and C.P. Research in the lab of J.-L.M. is supported by the Institut Curie, the Centre National de la Recherche Scientifique (CNRS), the Institut National de la Santé Et de la Recherche Médicale (INSERM), and is funded by grants from the Fondation Schlumberger pour l’Ēducation et la Recherche via the Fondation pour la Recherche Médicale, the European Molecular Biology Organization Young Investigator program (EMBO YIP), Labex DEEP (ANR- 11 -LABX-0044, part of the IDEX PSL ANR-10-IDEX-0001–02). J.F. is funded by a fellowship from the Fondation pour la Recherche Médicale (FDM202006011290). The work by H.T. and N.E. was supported by the CNRS and Collège de France. No fund from the European Research Council was used for this project.

## Author contributions

J.F., H.T., C.P. and J.-L.M. conceptualized the project and acquired funding. J.F. and J.-L.M. designed experiments. J.F. performed experiments. J.F. and J.-L.M. analyzed the data. J.F., D.N.D., V.B.L and C.P. organized embryo collection. N.E. and H.T. wrote the theory and performed numerical simulations. J.-L.M. wrote the manuscript with inputs from J.F., N.E., H.T. and C.P.

## Methods

### Ethics statement

The use of human embryos donated for this project was allowed by the Agence de la Biomédecine (ABM, approval number RE 17- 011R) in compliance with the International Society for Stem Cell Research guidelines^32^. All human preimplantation embryos used were donated after patients had fulfilled all reproductive needs. Informed written consent was obtained from both patients of all couples that donated embryos following IVF treatment. No financial incentive were offered for the donation.

Donated embryos were cryopreserved and stored at Fertilité Paris Centre ART Center (Biologie de la reproduction-CECOS, Cochin, APHP.Centre-Université de Paris), Clinique La Muette (Paris, France) or Clinique Pierre Chérest (Neuilly sur Seine, France). Embryos were then transferred to the Institut Curie where they were immediately thawed and used for the research project.

### Patients and embryos

A total number of 43 embryos provided by 33 couples of patients have been used for this study. Embryos were frozen on day 2 (n = 24) or day 3 (n = 19) according to slow freezing procedure (n = 30) or vitrification (n = 13). The mean cell number, at frozen time, was 4 ± 1 cells (mean ± SD, minimum 2 and maximum 8 cells) and 8 ± 1 cells (mean ± SD, minimum 6 and maximum 10 cells) for day 2 and day 3 frozen embryos respectively. For measurements throughout compaction (Fig 1, 3), day 2 embryos were thawed. For measurements on compacted embryos (Fig 2c-i) both day 2 and day 3 embryos were thawed. For immunostained embryos (Fig 2a-b), day 3 embryos were thawed.

Embryos were frozen for 11.2 ± 4.9 years (Mean ± SD, 13.6 ± 4.0 and 5.7 ± 1.1 years for slow-freeze and vitrified embryos respectively).

The donors mean ages were 34.1 ± 3.5 and 36.3 ± 6.3 year-old for female and male patients respectively (mean ± SD, data available for 30 of the 33 couples, data currently missing for 3 couples). 31/41 embryos were generated by intracytoplasmic sperm injection (ICSI) and 10/41 embryos were conceived using classical IVF. All sperms were fresh.

### Embryo work

#### Thawing

Embryos were handled using Stripper micropipettes (Origio) on binoculars (Leica M80) equipped with heating plates (Leica MATS-Type TL base) set to 37°C when needed.

Cryopreserved embryos were thawed according to manufacturer’s instructions. Embryo thawing packs (Origio) were used for slow-freeze embryos. Vit Kit (Irvine Scientific) was used for vitrified embryos. The intact survival rate, defined as the percentage of embryos without any cell lysis immediately after thawing, was 88,6% (39/43) for the embryos further considered for experimentation. All embryos survived the warming process with at least 50% of intact cells.

#### Culture

Embryos were handled using Stripper micropipettes (Origio) on binoculars (Leica M80) equipped with heating plates (Leica MATS-Type TL base) set to 37°C. Embryos are placed in pre-equilibrated (at least 4 h at 37°C, 5% O_2_, 5% CO_2_) CSCM-C medium (Irvine Scientific) covered with mineral oil (Irvine Scientific) in 50 mm glass bottom dishes (MatTek Corporation P50G-1,5-30-F) within an incubator (New Brunswick Galaxy 48 R) or the incubation chamber of the microscopes (Leica DMI6000 B with custom incubation from EMBLEM or Zeiss CellDiscoverer 7 with a 37°C humidified atmosphere supplemented with 5% CO_2_ and depleted to 5% O_2_ by supplementing N_2_.

#### Zona pellucida dissection

Before surface tension measurements, embryos were dissected out of their *zona pellucidae* (ZP) on the day of their thawing using a holding pipette and a glass needle^33^. The holding pipette and needle were custom-made from glass capillaries (Harvard apparatus, GC100TF-15) pulled using a P-97 Flaming Brown needle puller (Sutter Instrument) with the following settings: Ramp +25, Pull 65, Velocity 80, Time 175 and Pressure 200. To forge the holding pipette, the glass needle was cut to a ~120 μm diameter using a microforge (Narishige, MF-900) and the tip was fire-polished to a ~20 μm inner diameter. To forge the needle, the tip was melted onto the glass bead and pulled back to obtain a solid pointed tip. Both needle and pipette were bent to a 20° degrees angle to be parallel to the dish surface when mounted on the micromanipulator (Leica, AM6000). The holding pipette can apply controlled pressures using mineral oil-filled tubing coupled to a piston of which the position is moved using a microscale translating stage (Eppendorf, CellTram Oil 5176).

#### Chemical reagents

ML7 (Sigma-Aldrich, I2764) was diluted in 50% EtOH to 10 mM. Day 4 compacted embryos were placed into medium containing 1:2000 EtOH for 15 min before surface tension measurements for an additional 30 min. Embryos were then moved into medium containing 10 μM ML7 for 15 min before surface tension measurements taking another 30 min. Embryos were then placed back into normal culture medium CSCM-C to recover. 6/6 embryos recompacted and 5/6 embryos formed a blastocyst.

Similarly, day 4 compacted embryos were placed into Embryo Biopsie Medium (IrvineScientific), a commercial medium without Ca^2+^ and supplemented with 0.5 mM EDTA for 10 min before surface tension measurements taking another 20 min. Embryos were then placed back into normal culture medium CSCM to recover. 6/6 embryos recompacted and 4/6 embryos formed a blastocyst.

### Immunostaining

Embryos were fixed in 2% PFA (Euromedex, 2000-C) for 10 min at 37°C, washed in PBS and permeabilized in 0.01% Triton X-100 (Euromedex, T8787) in PBS (PBT) at room temperature before being placed in blocking solution (PBT with 3% BSA) at 4°C for at least 4 h. Primary antibodies were applied in blocking solution at 4°C overnight. After washes in PBT at room temperature, embryos were incubated with secondary antibodies, DAPI and phalloidin at room temperature for 1 h. Embryos were washed in PBT and imaged immediately after in PBS with BSA under mineral oil.

| Primary antibodies | Dilution | Provider | RRID |
| --- | --- | --- | --- |
| MYH10 | 1:200 | Santa Cruz, sc-376942 |  |
| CDH1 | 1:200 | Cell Signaling Technology, 3195 | AB_2291471 |

| Secondary antibodies and dyes | Dilution | Provider | RRID |
| --- | --- | --- | --- |
| Alexa Fluor Plus 488 anti-mouse | 1:200 | Invitrogen, A32723 | AB_2633275 |
| Alexa Fluor 546 anti-mouse | 1:200 | Invitrogen, A11003 | AB_2534071 |
| Alexa Fluor Plus 488 anti-rabbit | 1:200 | Invitrogen, A32731 | AB_2633280 |
| Alexa Fluor Plus 546 anti-rabbit | 1:200 | Invitrogen, A11010 | AB_2534077 |
| Alexa Fluor 633 phalloidin | 1:200 | Invitrogen, A22284 |  |
| 4',6-diamidino-2-phenylindole (DAPI) | 1:1000 | Invitrogen, D1306 | AB_2629482 |

### Micropipette aspiration

#### Micropipette preparation

To forge the micropipettes, glass capillaries (World Precision Instruments TW100-3) were pulled using a P-97 Flaming Brown needle puller (Sutter Instrument) with the following settings: Ramp +3–5, Pull 55, Velocity 50, Time 50 and Pressure 500.

Using a microforge (Narishige, MF-900), needles were cut to form a blunt opening of radius 12-22 μm and bent 80-100 μm away from the tip at a 20° angle.

#### Microaspiration setup

The micropipette was mounted on a micromanipulator (Leica AM6000) using a grip head and capillary holder (Eppendorf, 920007392 and 9200077414). The micropipette was connected to a PBS-filled intermediate reservoir of which the height is controlled using a 50 mm microscale translating stage (Newport) to generate positive and negative pressures^34^. The intermediate reservoir was connected to a microfluidic pump (Fluigent, MFCS-EZ) delivering negative pressures with a 2.5 Pa resolution. The pressure is controlled using Maesflow software (Fluigent). The output pressure was calibrated by finding the height of the intermediate reservoir at which no flow in the micropipette is observed (using floating particles found in the dish, ‘no flow’ is considered achieved when the position of the particle inside the micropipette is stable for ~10 s and, if slow drift can be reverted with 10 Pa).

#### Surface tension measurement

To measure cell surface tension, the micropipette was brought in contact with the free surface of a blastomere of an embryo with a low grabbing pressure (20-30 Pa, Fig. 1a). The pressure was then increased stepwise (10 Pa steps) until the deformation of the blastomeres reaches the radius of the micropipette (*R_p_*). At steady state, the surface tension *γ*_cm_ of the blastomeres was calculated using Young–Laplace’s law: *γ*_cm_ = *P_c_* / 2 (1/*R_p_* - 1/*R_c_*), where *P_c_* is the pressure used to deform the cell of radius *R_c_*. The pressure was then released and relaxation of the deformation was observed. It typically took 3-5 min to probe a cell.

#### Interfacial tension measurement

After measuring the surface tension of two adjacent cells of the embryo, we assumed steady state on the timescale of the measurement and calculate the interfacial tension from the force balance equation at the cell-cell contact (Fig 1a). On the basis of the general Young-Dupré equation, we calculated *γ*_cc_ = - *γ*_cm1_ cos (θ_1_) - *γ*_cm2_ cos (θ_2_), where *γ*_cm1_ and *γ*_cm2_ are the surface tensions of cell 1 and 2, θ1 and θ_2_, the internal contact angles of cell 1 and 2, and *γ*_cc_, the interfacial tension at the cell-cell contact. In the approximation of contacting cells with equivalent surface tensions *γ*_cm_, the contribution of each blastomereto *γ*_cc_ is *γ*_cc_/2^20^.

### Microscopy

#### Pipette-scope

Surface tension measurements were performed on a Leica DMI6000 B inverted microscope equipped with a 40x/0.8 DRY HC PL APO Ph2 (11506383) objective. A 0.7x lens is mounted in front of a Retina R3 camera. The microscope is equipped with a custom incubation chamber (EMBLEM) to keep the sample at 37°C and maintain the atmosphere at 5% CO_2_ and 5% O_2_.

#### Time lapse imaging

For time-lapse imaging, embryos were placed after thawing within the chamber of a CellDiscoverer 7 (Zeiss) humidified 37°C, 5% O_2_, 5% CO_2_ atmosphere. Embryos were imaged every 30 min at 5 to 8 focal planes separated by 10 *μ*m using IR-LED (725 nm) transmitted light through a 20x/0.95 objective. Images were acquired using either an ORCA-Flash 4.0 camera (Hamamatsu, C11440) or a 506 axiovert (Zeiss) camera.

#### Spinning disc microscope

Immunostainings were imaged on a Zeiss Observer Z1 microscope with a CSU-X1 spinning disc unit (Yokogawa) using 405, 488, 561, and 642 nm laser lines through a 63x/1.2 C Apo Korr water immersion objective; emission was collected through 450/ 50 nm, 525/50 nm, 595/50 band pass or 610 nm low pass filters onto an ORCA-Flash 4.0 camera (C11440, Hamamatsu).

### Data analyses

#### Image analysis

Pipette size, cell radii of curvature, contact angles were measured using Fiji with the line, circle and angle tools respectively^35^.

To measure cortical intensity, as done previously^20^, we picked confocal slices cutting through the equatorial plane of two contacting cells using Fiji. We drew a 1 μm thick line along the cell-medium interface or cell-cell contact, measured the mean intensity and divided the contact intensity by the sum of the cell-medium intensities of contacting cells.

#### Statistics

Mean, standard deviation, SEM, Pearson’s correlation coefficients, were calculated using Excel (Microsoft). Statistical tests were performed with the free online tool BiostaTGV (https://biostatgv.sentiweb.fr/), based on R. Two-tailed Student t-tests and Pearson’s correlation tests were performed when needed. Statistical significance was considered when *p* < 10^-2^.

Contact angle thresholds were determined using Cutoff Finder^26^, a bundle of optimization and visualization methods for cutoff determination based on R (https://molpathoheidelberg.shinyapps.io/CutoffFinder_v1/).

Non-compacting embryos and fully compacting embryos were qualitatively assessed according to ESHRE guidelines^36^. Maximizing specificity (100%) for detecting non-compacting embryos gave a sensitivity of 82% and a cutoff angle at 131° with an area under the curve (AUC) of 0.991. For partially compacting embryos, qualitative assessment, following recent descriptions of the phenomenon^4,15^, yielded a cutoff angle of 113° is found with 100% specificity, 100% sensitivity and AUC of 1.

The sample size was not predetermined and simply results from the repetition of experiments. No sample was excluded. No randomization method was used. The investigators were not blinded during experiments.

## Data and code availability

Images and data used for the analysis will be made available on a public repository.

Code is available upon request.

## Supplementary material

**Extended Data Figure 1:**
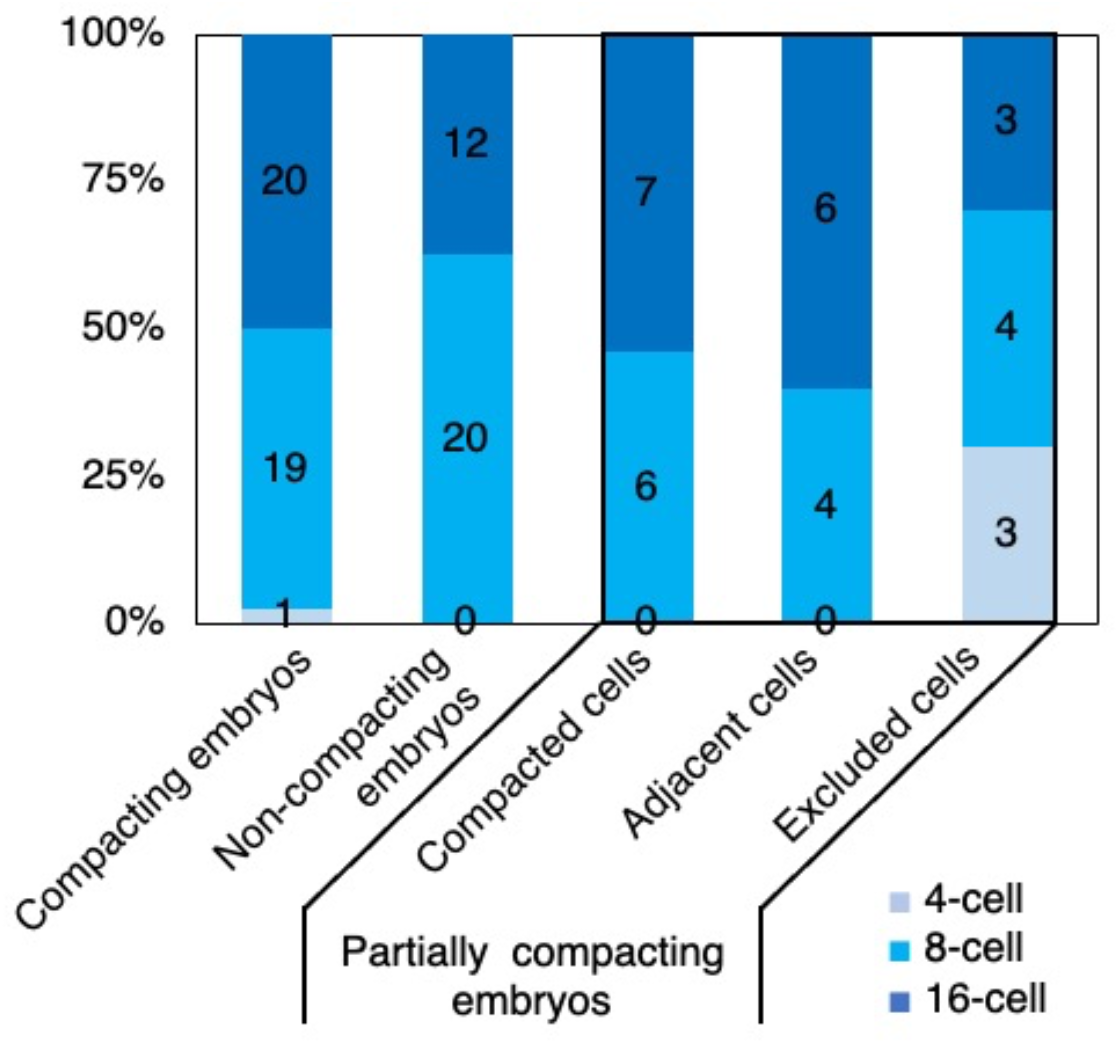
cleavage stage of compacting and non-compacting blastomeres. Blastomere stage, as determined by tracking successive cell divisions until the last tension measurement, of fully compacting, non-compacting and partially compacting embryos (7, 6 and 6 embryos and 40, 32 and 33 blastomeres respectively). For partially compacting embryos, the stages of compacted blastomeres, compacted blastomeres adjacent to excluded cells and excluded cells are indicated separately (13, 10 and 10 respectively).

**Movie 1: preimplantation development of human embryos with full compaction.** Time-lapse imaging of a human embryo from the 4-cell stage to the blastocyst stage (shown in Fig 1b). Pictures taken every 30 min, scale bar 40 *μ*m.

**Movie 2: preimplantation development of human embryos without compaction.** Time-lapse imaging of a human embryo from the 4-cell stage to the 16-cell stage (shown in Fig 3a). Pictures taken every 30 min, scale bar 40 *μ*m.

**Movie 3: preimplantation development of human embryos with partial compaction.** Timelapse imaging of a human embryo from the 4-cell stage to the blastocyst stage (shown in Fig 3e). Pictures taken every 30 min, scale bar 40 *μ*m.

**Movie 4: 3D simulations of compaction.** Simulations of compaction with distinct cell populations: blue blastomeres grow their tension *γ*_cm_ by a factor 3.2 and their tension *γ*_cc_ by 1.4 according to measurements shown in Fig 3g-h; purple blastomeres do the same as blue ones (bottom left), keep *γ*_cm_ steady and grow their tension *γ*_cc_ by 1.4 (bottom middle) or keep *γ*_cm_ steady and grow their tension *γ*_cc_ by 2.4. Tensions are linearly interpolated between the initial and final states in 15 steps.

**Supplementary table 1:**
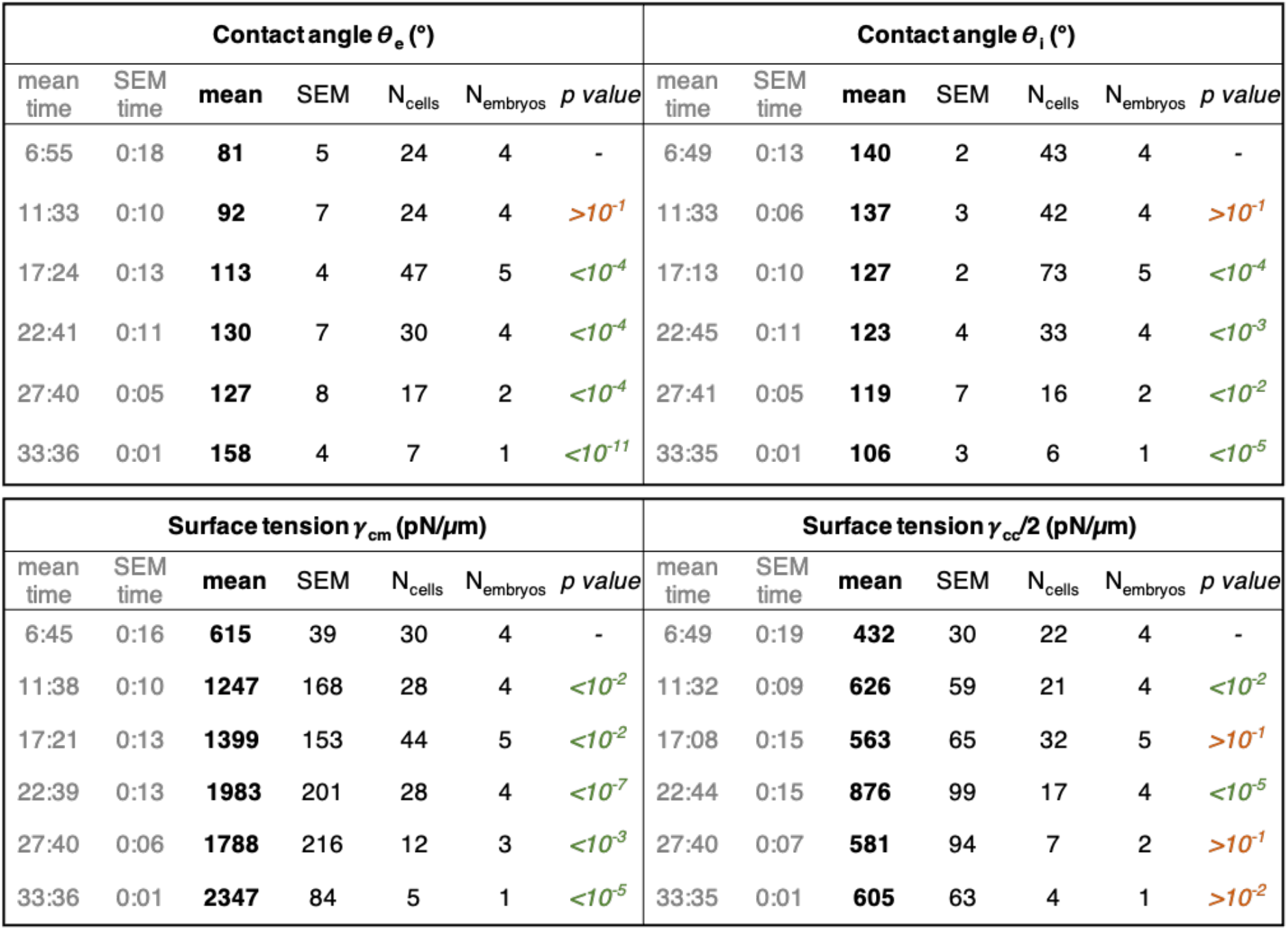
Contact angles and surface tensions of compacting embryos synchronized to the time of last observed 3^rd^ cleavage. Mean, standard error of the mean (SEM) and numbers of cells and embryos for the time after last observed 3^rd^ cleavage, external contact angle θ_e_, internal contact angle θ_i_, surface tensions *γ*_cm_ and *γ*_cc_/2. Data binned over 5 h, as shown in Fig 1c-d. p values given for Student’s t test between a given time point and the first one for the contact angles and surface tensions. p values in green when below 10^-2^ and in red when above.

**Supplementary table 2:** Contact angles and surface tensions of compacted embryos before and after treatment with ML7 or EDTA media. Mean, standard error of the mean (SEM) and numbers of cells and embryos for the external contact angle θ_e_, internal contact angle θ_i_, surface tensions *γ*_cm_ and *γ*_i_. Data shown in Fig 2d-e and 2g-h. p values given for pairwise Student’s t test between the control and treatment. p values in green when below 10^-2^ and in red when above.

| | Contact angle $\theta_e$ (°) | | | | | Contact angle $\theta_i$ (°) | | | | |
| --- | --- | --- | --- | --- | --- | --- | --- | --- | --- | --- |
|  | mean | SEM | N<sub>cells</sub> | N<sub>embryos</sub> | p value | mean | SEM | N<sub>cells</sub> | N<sub>embryos</sub> | p value |
| **Ctrl** | **133** | 5 | 59 | 6 | - | **114** | 3 | 80 | 6 | - |
| **ML7** | **106** | 4 | 51 | 6 | $<10^{-4}$ | **126** | 2 | 77 | 6 | $<10^{-2}$ |

| | Surface tension $\gamma_{cm}$ (pN/μm) | | | | | Surface tension $\gamma_{cc}/2$ (pN/μm) | | | | |
| --- | --- | --- | --- | --- | --- | --- | --- | --- | --- | --- |
|  | mean | SEM | N<sub>cells</sub> | N<sub>embryos</sub> | p value | mean | SEM | N<sub>cells</sub> | N<sub>embryos</sub> | p value |
| **Ctrl** | **1482** | 221 | 55 | 6 | - | **598** | 74 | 37 | 6 | - |
| **ML7** | **514** | 91 | 60 | 6 | $<10^{-2}$ | **271** | 53 | 79 | 6 | $<10^{-3}$ |

| | Contact angle $\theta_e$ (°) | | | | | Contact angle $\theta_i$ (°) | | | | |
| --- | --- | --- | --- | --- | --- | --- | --- | --- | --- | --- |
|  | mean | SEM | N<sub>cells</sub> | N<sub>embryos</sub> | p value | mean | SEM | N<sub>cells</sub> | N<sub>embryos</sub> | p value |
| **Ctrl** | **141** | 6 | 54 | 6 | - | **109** | 2 | 80 | 6 | - |
| **EDTA** | **88** | 5 | 45 | 6 | $<10^{-2}$ | **137** | 2 | 69 | 6 | $<10^{-3}$ |

| | Surface tension $\gamma_{cm}$ (pN/μm) | | | | | Surface tension $\gamma_{cc}/2$ (pN/μm) | | | | |
| --- | --- | --- | --- | --- | --- | --- | --- | --- | --- | --- |
|  | mean | SEM | N<sub>cells</sub> | N<sub>embryos</sub> | p value | mean | SEM | N<sub>cells</sub> | N<sub>embryos</sub> | p value |
| **Ctrl** | **1361** | 192 | 49 | 6 | - | **396** | 81 | 35 | 6 | - |
| **EDTA** | **1439** | 211 | 49 | 6 | $>10^{-1}$ | **1048** | 167 | 26 | 6 | $<10^{-2}$ |

**Supplementary table 3:**
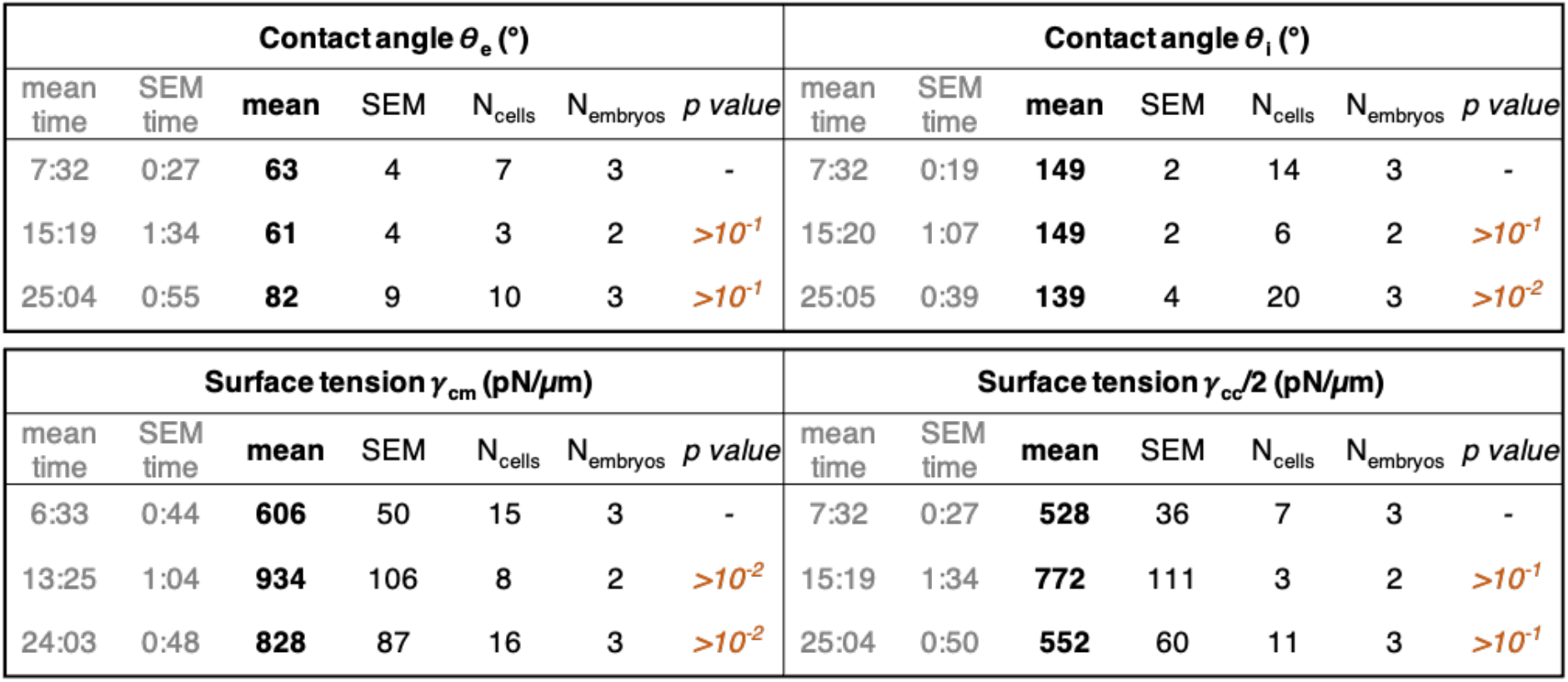
Contact angles and surface tensions of non-compacting embryos synchronized to the time of last observed 3^rd^ cleavage. Mean, standard error of the mean (SEM) and numbers of cells and embryos for the time after last observed 3^rd^ cleavage, external contact angle θ_e_, internal contact angle θ_i_, surface tensions *γ*_cm_ and *γ*_cc_/2. Data binned over 10 h, as shown in Fig 3b. p values given for Student’s t test between a given time point and the first one for the contact angles and surface tensions. p values in green when below 10^-2^ and in red when above.

**Supplementary table 4:**
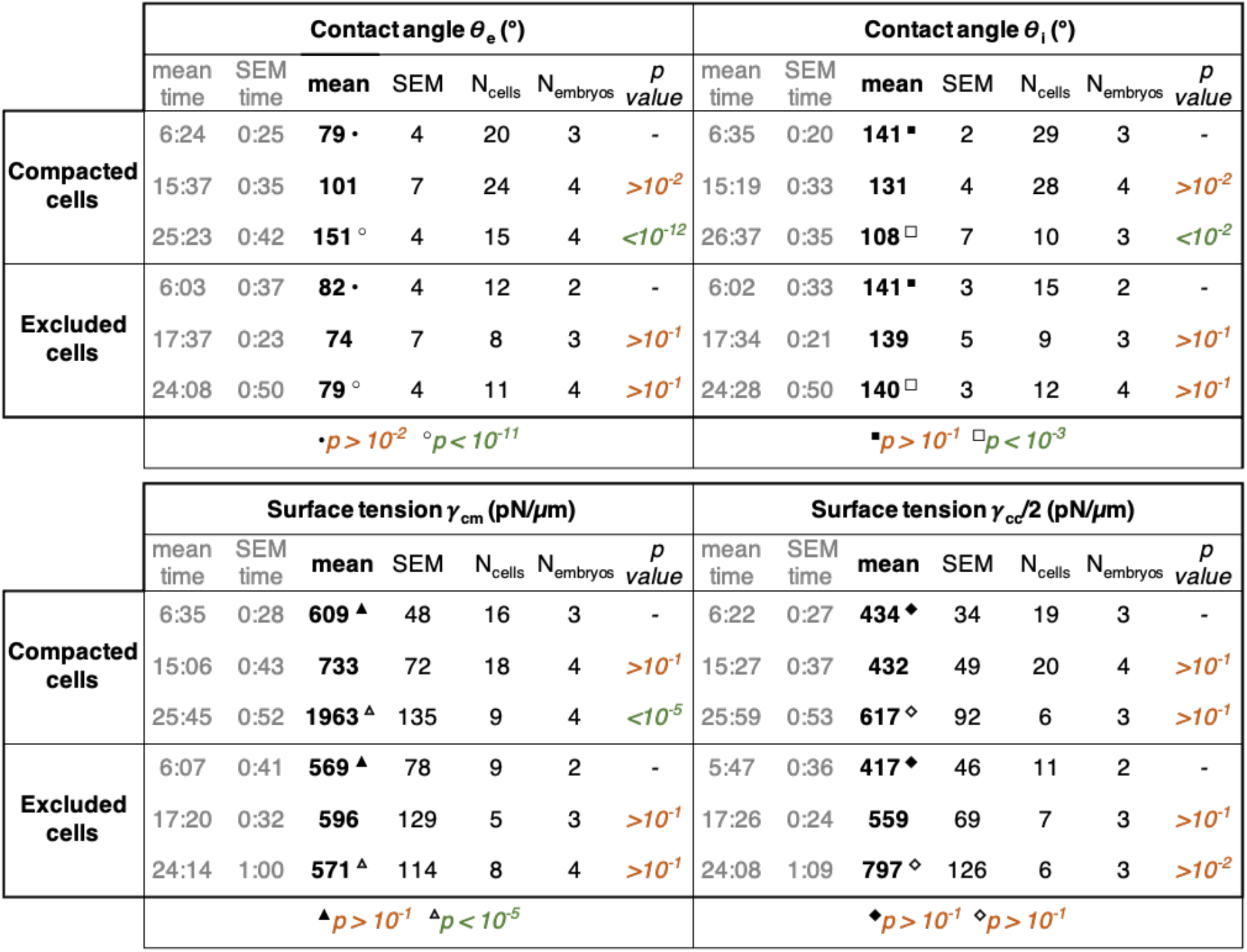
Contact angles and surface tensions of partially compacting embryos synchronized to the time of last observed 3^rd^ cleavage. Mean, standard error of the mean (SEM) and numbers of cells and embryos for the time after last observed 3^rd^ cleavage, external contact angle θ_e_, internal contact angle θ_i_, surface tensions *γ*_cm_ and *γ*_cc_/2. Data binned over 10 h for compacted and excluded cells as shown in Fig 3f-h. p values given for Student’s t test between a given time point and the first one for the contact angles and surface tensions. p values in green when below 10^-2^ and in red when above.

## SUPPLEMENTARY NOTE

3D active foam model

In this Supplementary Note, we detail the assumptions of our physical model, its numerical implementation and the choice of simulation parameters.

### 1 Model hypotheses

In our study, we assume that the shape of blastomeres in the preimplantation human embryo is controlled primarily by cortical tensions, in a similar manner as in other early embryos (1–3). These surface tensions originate from cortical contractility and are supposed homogeneous and isotropic at each cell-cell interface. The tension of the plasma membrane is generically lower by an order of magnitude (3) and the direct negative contribution to the surface tension of adhesion molecules (cadherin) is generally negligible as well (2). An assembly of cells is therefore akin to a foam, but where each interface may have a different surface tension, that is controlled actively by adjacent cells. The evolution of cell shapes is assumed to be quasistatic, which means that viscous dissipation may be neglected. This hypothesis is well-justified from the comparison of the typical timescale associated to viscous relaxation of the cortex, of the order of minutes (4, 5), and the timescale associate with compaction, of the order of tens of hours. Moreover, we assume that blastomeres conserve their volumes during the compaction process, as previously measured in mouse embryos (2).

### 2 Cell doublet: a toy model for compaction

For a cell doublet, we recall below some results presented in (2) and (3). The doublet’s shape can be parametrized by the two cell volumes *V*_1_ and *V*_2_ and three surface tensions: *γ*_cm1_ and *γ*_cm2_ at the cellmedium interfaces of cell 1 and 2, and *γ*_cc_ at the cell-cell contact. Force balance in the doublet may be summarized by four equations

- Laplace’s law in cell 1: 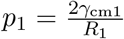, where *R*_1_ is the curvature radius of the cell-medium interface of cell 1.
- Laplace’s law in cell 2: 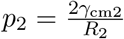, where *R*_2_ is the curvature radius of the cell-medium interface of cell 2.
- Laplace’s law at the cell-cell contact: 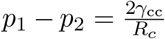, where *R_c_* is the curvature radius of the contact interface between the two cells.
- Young-Dupré’s law at the cell-cell junction: 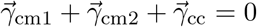,

which set the geometry of the doublet, together with volume constraints 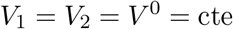.

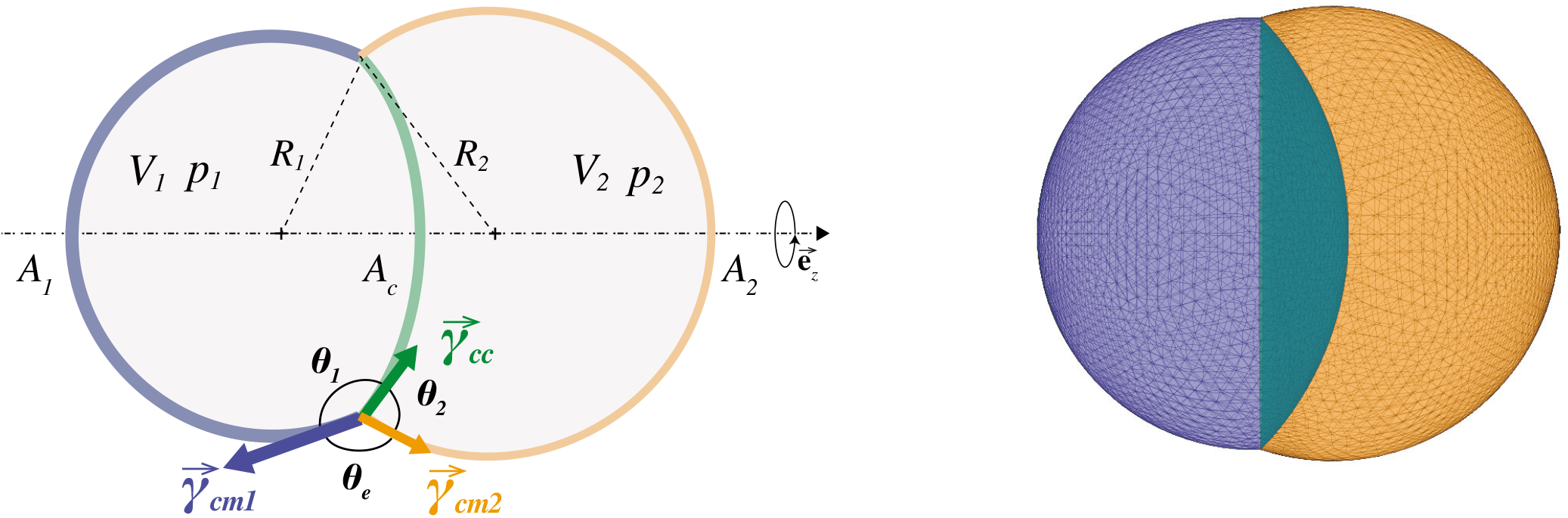
(Left) Schematic representation of a cell doublet, with surface tensions at cell-medium and contact interfaces *γ*_cm1_, *γ*_cm2_ and *γ*_cc_, surface areas *A*_1,2_, interface area *A_c_*, cell volumes *V*_1,2_, pressures *p*_1,2_, curvature radii *R*_1,2_, and contact angles *θ*_1,2_ and *θ_e_.* The curvature radius of the contact interface *A_c_* is called *R_c_*. (Right) Corresponding simulation triangular mesh.

The law of Young-Dupré may be rewritten as function of internal and external contact angles *θ*_1_, *θ*_2_ and *θ_e_* as follows:

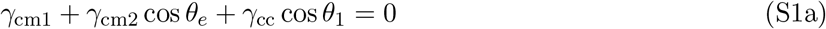

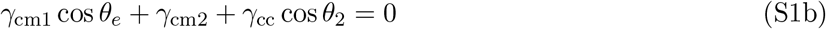

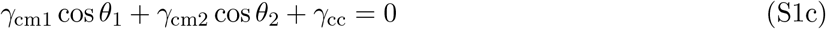

One may note that the above force balance equations are not restricted to cell doublets, but are valid at any contacts and junctions between cells. In the next, we therefore apply them directly to human embryos.

#### Compaction parameter

By combining the Young-Dupré laws in Section 2 as follows Eq. S1a\**γ*_cm1_ + Eq. S1b\**γ*_cm2_ - Eq. S1c\**γ*_cc_ leads to

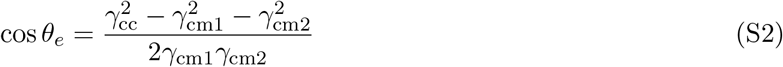

In the case of a symmetric doublet, or symmetric cell-cell contact, i.e. when *γ*_cm1_ = *γ*_cm2_ = *γ*_cm_, the equation Eq. S2 simplifies into

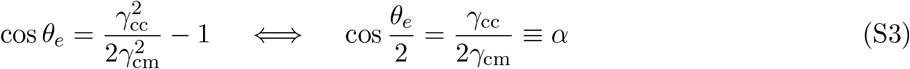

where we used the trigonometric relation cos 2*a* = cos^2^ *a*–sin^2^*a*. The parameter α was named ***compaction parameter*** in earlier studies (2, 6) and varies between 0 and 1. Increasing compaction corresponds to a decreasing parameter *α* as illustrated on the phase diagrams Figs. 1g and 2i in the main text.

#### Minimal compaction path

Below we define the notion of *minimal* path in the parameter space of our phase diagrams in Figs. 1g and 2i, where the time course of compaction is plotted against *γ*_cm_ in x-axis and *γ*_cc_/2 in y-axis. We first note that this idea of “minimal” here is not related to any consideration of cell metabolism or mechanical dissipation and is not meant to imply an evolutionary advantage. It simply describes the path that requires smallest changes in the individual cell surface tensions *γ*_cm_ and *γ*_cc_/2 to reach a given state of compaction f starting from an initial configuration *i*. The initial configuration is characterized by two given tension values 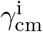 and 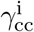, leading to a compaction parameter denoted 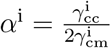 at initial state,. The final state is only characterized by the final compaction parameter α^f^.

The problem may be reformulated as a geometric problem, where we look for the minimal distance between a point 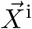 of coordinates 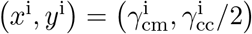 and the line of slope *y/x* = α^f^ going through the origin. This minimal distance is obtained as the unique intersection point 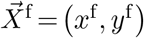 between the line of slope α^f^ and the perpendicular line going through the point 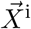. Mathematically, this may be formulated as below

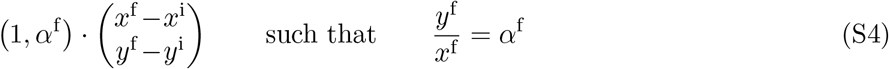

the solution of which is given by

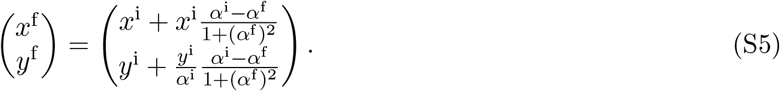

Denoting by Δ*x* = *x*^f^ – *x*^i^ and Δ*y* = *y*^f^ – *y*^i^ absolute changes in surface tensions, we deduce immediately the *minimal* tensions fold changes necessary to reach the final compaction state as

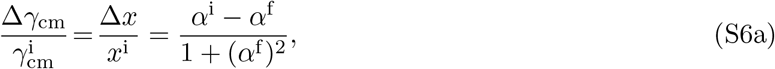

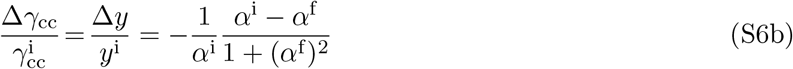

In human embryos, the compaction parameter *α* decreases from 0.70 to 0.26 by increasing *γ*_cm_ by 282% and increasing *γ*_cc_ by 40%. This is in sharp contrast with the fold changes predicted for the minimal path between these compaction states, where *γ*_cm_ would increase by 11% and *γ*_cc_ would decrease by 59%.

In comparison, in the mouse, the compaction parameter α decreases from 0.72 to 0.28 by increasing *γ*_cm_ by 97% and decreasing *γ*_cc_ by 24.4%. This path is closer to the predicted minimal one, where *γ*_cm_ would have to increase by 11.3% and *γ*_cc_ to decrease by 56.9%.

Graphically, these minimal paths can be plotted on the phase diagram in log-log scale. To allow for comparison, between mouse and human cases, we have normalized in the figure below all tensions by their initial values *γ*_cm0_ or *γ*_cc0_.

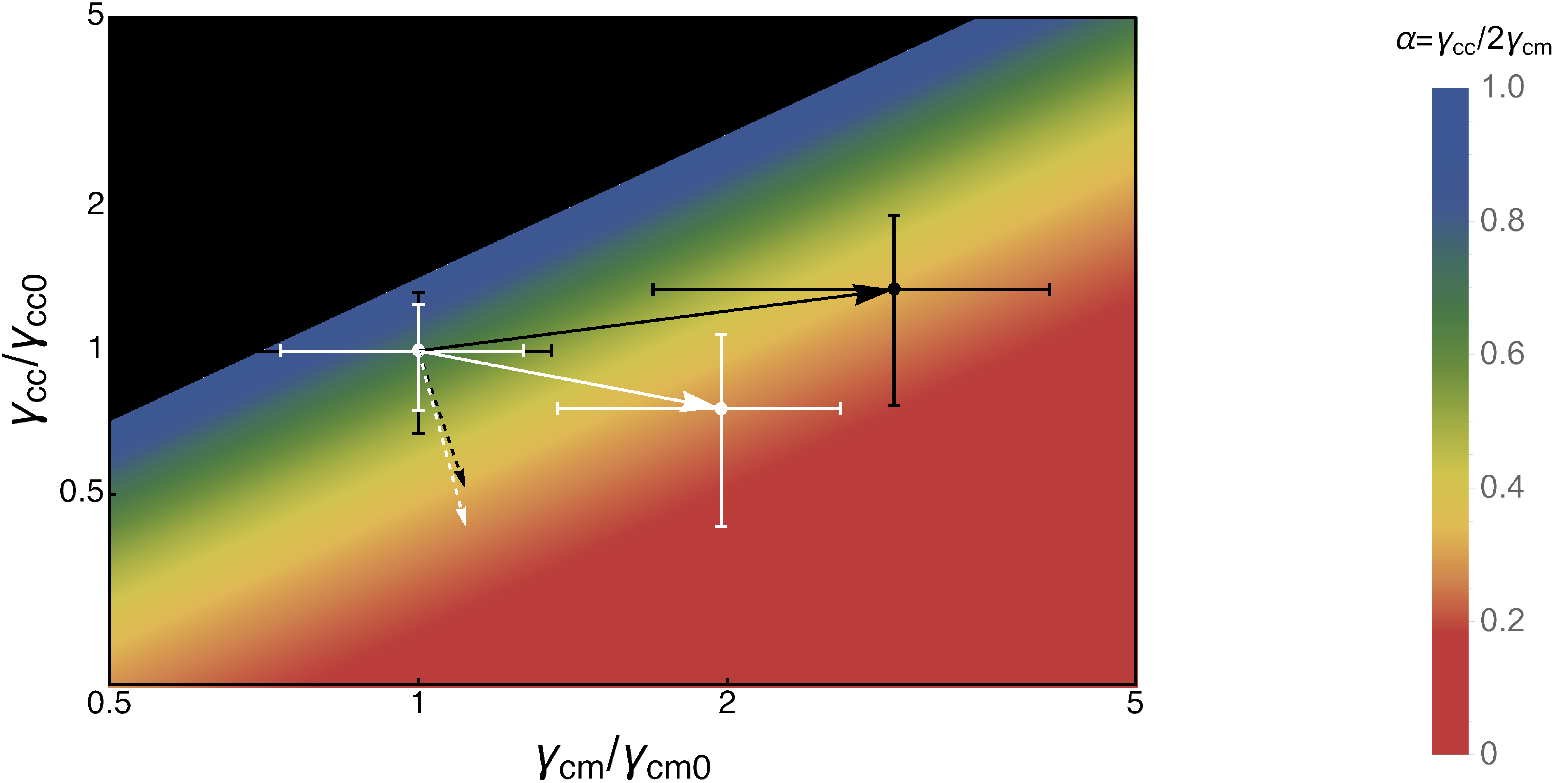
Normalized tension phase diagram in log-log plot, comparing the experimental evolution of compaction in human and mouse embryos (plain lines in black and white colors respectively) with corresponding ones that would minimize tension changes (dashed lines). The compaction state 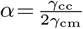 is color-coded for better visualization purposes.

This plot shows that both mouse and human embryos are far from the minimal compaction path in such phase diagram, but that the mouse embryo performs better in minimizing its tensions changes to reach its final compaction state.

#### Surface energy and Lagrangian function

It was shown previously in reference (3), that the shape of the doublet may be also obtained by minimizing a surface energy 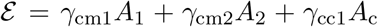, where *A*_1_, *A*_2_ and *A_c_* are respectively the area of the cell-medium interfaces of cells 1 and 2 and of the cell-cell contact, while conserving cell volumes. Such constrained optimization problem may be solved practically through the introduction of a Lagrangian function, where the two Lagrange multipliers *p*_1_ and *p*_2_ are in fact the cell pressures

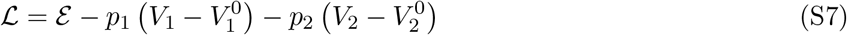

where 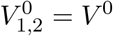 are the target volumes values.

Optimality conditions are obtained by setting to zero the derivatives of the Lagrangian function with respect to the doublet shape geometry and Lagrange multipliers (7). Below, we exemplify this approach by parametrizing the shape of cells with triangular meshes.

### 3 Simulations with *N* cells

#### 3.1 Lagrangian function and force on a vertex

Generalizing the approach for a cell doublet above, our numerical simulations consist in a constrained optimization of a surface energy defined on an initial non-manifold triangular mesh. The surface energy and Lagrangian function for a set of *N* cells are defined as follows

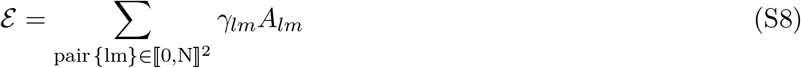

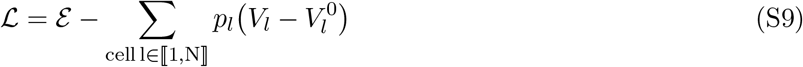

where *γ_lm_* and *A_lm_* are respectively the surface tension and area of the interface between regions *l* and *m,* and *p_l_* and 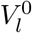 are respectively the pressure and the target volume value of the cell *l*. Note that for interfaces *l* and *m* span 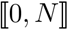, where 0 refers to the external medium, while for cells *l* spans 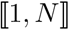.

From the Lagrangian function, one can calculate the force **f**_*k*_ on each vertex of the mesh 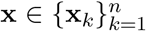 as follows

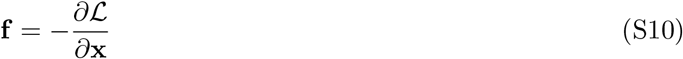

where **x** = *x***e**_*x*_ + *y***e**_*y*_ + *z***e**_*z*_ and **f**_*k*_ = *f*^*x*^**e**_*x*_ + *f*^*y*^**e**_*y*_ + *f*^*z*^**e**_*z*_ are the decomposition of vertex position and force in the 3D Euclidean space, equipped with an orthonormal basis (**e**_*x*_, **e**_*y*_, **e**_*z*_).

Interfacial areas and cell volumes can be easily expressed as sums on the triangles *t* in the mesh.

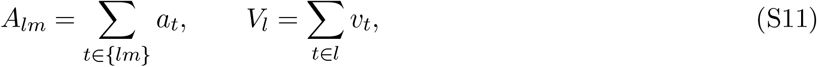

where *a_t_* and *v_t_* are respectively the elementary area and volume of a given triangle 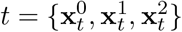, which are given by

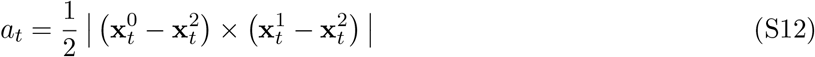

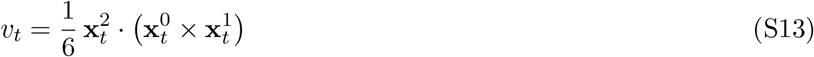

Their derivatives with respect to the vertex position 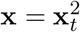 may be easily calculated as

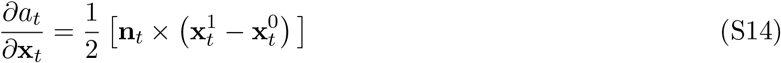

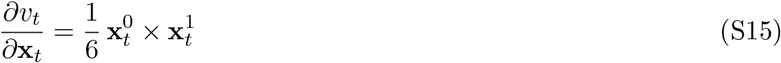

where we have defined the normal 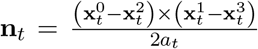 to the triangle *t*. Note that these formula are invariant by permutation of the triplet of vertices {0,1, 2}.

The force on a vertex **x** defined in Eq. S10 may now be explicitly expressed as

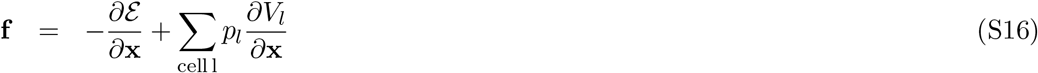

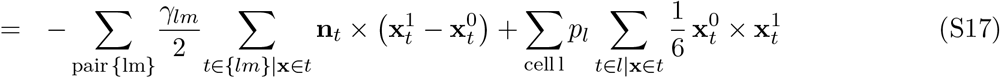

where we assumed without loss of generality that 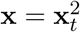, relying on the above invariance by permutation of derivatives formula.

#### 3.2 Constrained optimization: conjugate gradient and projection method

At mechanical equilibrium, all interfaces follow Laplace’s law and each junction verifies Young-Dupré’s equations. These equations may be equivalently expressed through a constrained optimization of the surface energy Eq. S8, where cell volumes are conserved. Using the Lagrangian function defined above in Eq. S9, optimality conditions (7) are obtained when

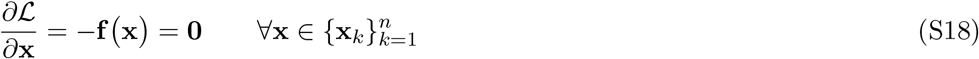

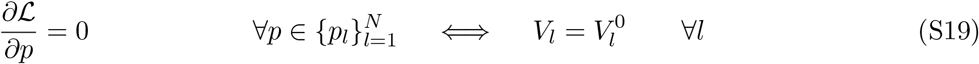

To calculate the Lagrange multipliers 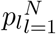, which enforce the volume conservation equations Eq. S19, we use a projection method. The force **f**_*k*_ = **f**(**x**_*k*_) of each vertex **x**_*k*_ is projected onto a subspace that is orthogonal to the space of cell volumes variations

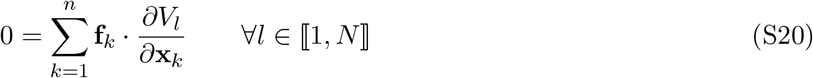

This leads to a linear system of equations for *p_l_*

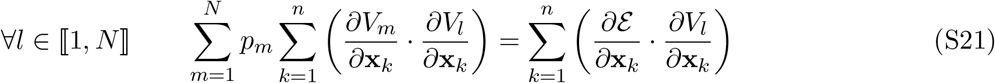

that may be rewritten in a condensed form 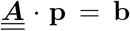, where **p** ≡ (*p*_1_,*p*_2_,⋯,*p_N_*) is a vector of size *N* collecting the unknown pressures, 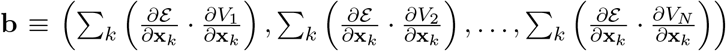 is the vector of constants and 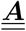 is the matrix of coefficients defined by

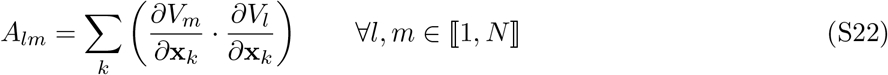

which is a symmetric positive definite and therefore invertible matrix. This linear system of *N* equations is solved using a Newton’s method (7).

To find the mechanical equilibrium, one needs to solve the equation Eq. S18: ∀**x**, **f**(**x**) = 0. To do so, we use an iterative method, namely the conjugate gradient, whereby the position **x**(*t*) of each vertex at time *t* is updated iteratively along a new search direction **D**_*t*+1_ that is conjugate to the one at previous time step **D**_*t*_ (7). Here, we follow the Hestenes-Stiefel conjugate-gradient scheme (8)

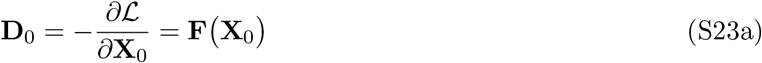

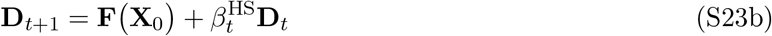

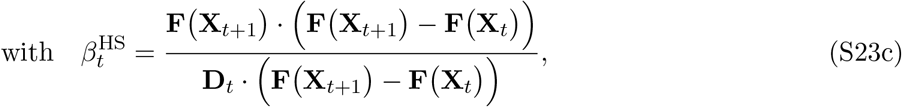

where we have defined a generalized vertex position vector **X**_*t*_ = **(x**_0_(*t*), **x**_1_(*t*),…, ***x***_*n*_(*t*) and a generalized force vector **F**(**X**_*t*_) = **f** (**x**_0_(*t*)), **f** (**x**_*n*_(*t*)),…, **f** (**x**_*n*_(*t*))).

At each iteration, the cell pressures are recalculated using the projection method as described above.

The constrained optimization method is applied to a non-manifold multimaterial mesh following (9). The identity of each interface separating cells *i* and *j* is tracked over its evolution by a label of integers *(i, j*) that is stored in each triangle of the interface. To maintain numerical precision, the triangular mesh is furthermore allowed to vary the number of vertices, edges and faces over its evolution (remeshing), and to perform topological transitions: T1 (neighbor exchange), T2 (region collapse) and merging (new contact). Note that, in contrast to classical vertex models where cell-cell boundaries are assumed to remain flat, our 3-dimensional model does not impose any prior constraints on cell shapes. Its precision to represent smooth and continuous interfaces is only limited by the user-defined resolution of the triangular mesh.

#### 3.3 Controls and choice of parameters

##### Controls for simulations of cell exclusion

For compaction, controls were already performed in (3) for simulations performed with the same numerical scheme by comparing them with semi-analytical results. Here, we therefore focus on the process of cell exclusion.

Following Eq. S2, a given set of tensions *γ*_cm1_ and *γ*_cm2_, the external contact angle *θ_e_* is related to the tension at the contact *γ*_cc_ as follows

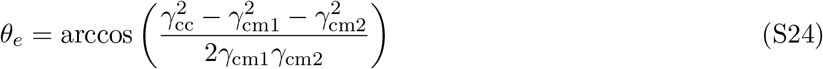

Choosing the two cell-medium tensions *γ*_cm1_ = 1.963 and *γ*_cm2_ = 0.609 for normal and excluded cells respectively, we can plot the external contact angle *θ_e_* at contacts with excluded cells as function of the value of the contact tension *γ*_cc_ using the above relation. Concurrently, we can perform simulations with corresponding values. Plotting the two against each others in the figure below, shows a very good agreement between simulations and theoretical predictions.

##### Choice of simulation parameters

We performed three simulations with *N* = 16 cells:

1. One where the embryo fully compacts, with initial and final values of the cell-medium surface tension *γ*_cm_ and contact tension *γ*_cc_ are the ones measured experimentally (Main text Fig. 3g,h). Note that experimentally *γ*_cc_ remains relatively constant between the initial (0h) and final state (30h).
2. One where one cell in this embryo has its cell-medium tension *γ*_cm_ maintained constant at its initial value as in the experiment, and *γ*_cc_ does the same.
3. A last one, where the tension *γ*_cm_ of one cell is still maintained fixed but the tension *γ*_cc_ at contacts with this cell is increased by a factor 2.4 from its initial value. This multiplicative factor is determined from the expected mean contact angle measured experimentally *θ_e_* ~ 78° for excluded cells after embryo compaction, using the relation Eq. S2 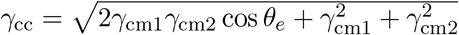, where 1 and 2 stand here for excluded and adjacent cells.

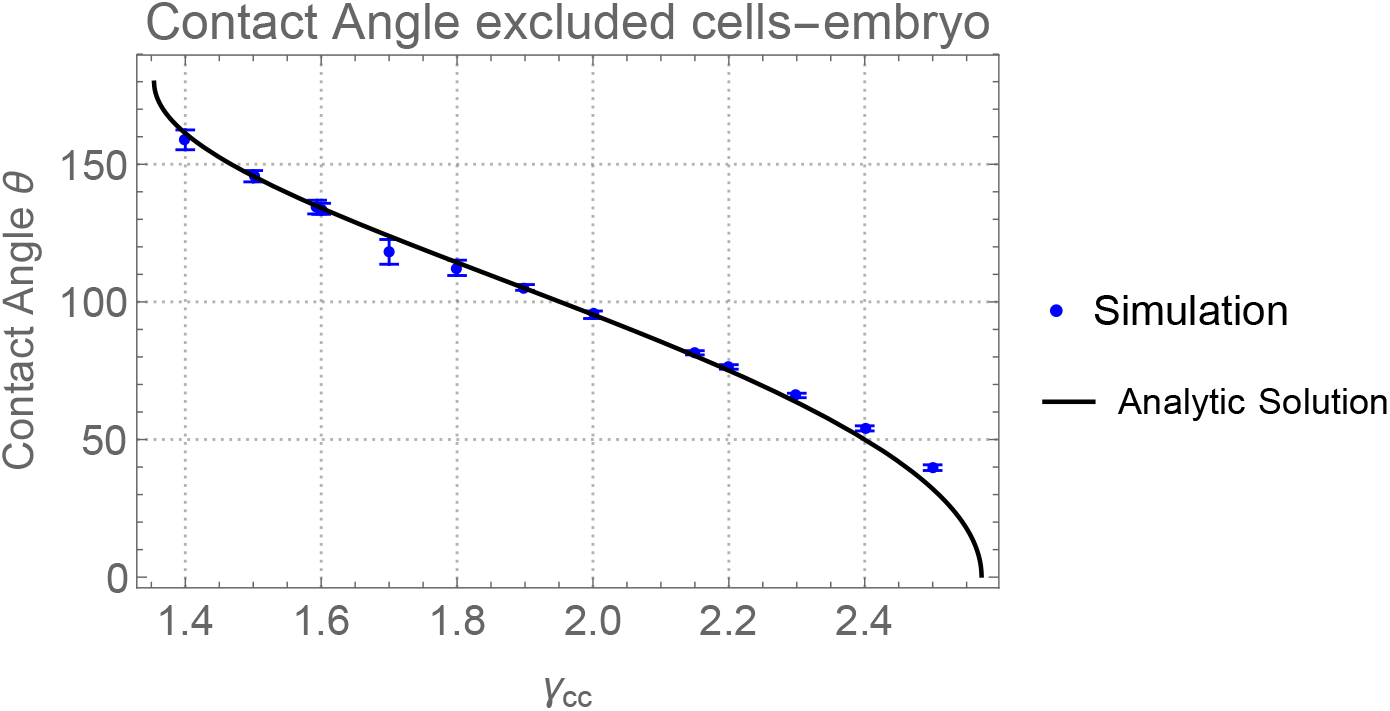
Plot of the contact angle between excluded and compacting cells as function of the value of the contact tension *γ*_cc_ at these contacts, for *γ*_cm2_ = 1.963 and *γ*_cm2_ = 0.609. The blue dots correspond to numerical simulations (mean and STD on the contact angles of 2 excluded cells with all of their individual neighbors), and the plain line corresponds to the the analytic formula Eq. S24.

To obtain a smooth evolution of cells shape, tension values are linearly interpolated in a total of 15 points from initial and final values. The initial and final tension values used in simulations are divided by a factor 1000 compared to experimental ones and separated by a right arrow in the Tables S1 below. A value in the column of label *i* and the line of label *j* corresponds to the interface surface tension between all cells {*ij*}, with *i, j* ∈ {0,.., 16}. Note that the label 0 corresponds to the external medium, such that values at the crossroad of 0 and a label *i* ≥ 1 give the cell-medium surface tension *γ_cm,i_* of a cell *i*. The tables are symmetric by definition of cell-medium and cell-cell interfaces *γ_ij_* = *γ_ji_*.

**Table S1:**
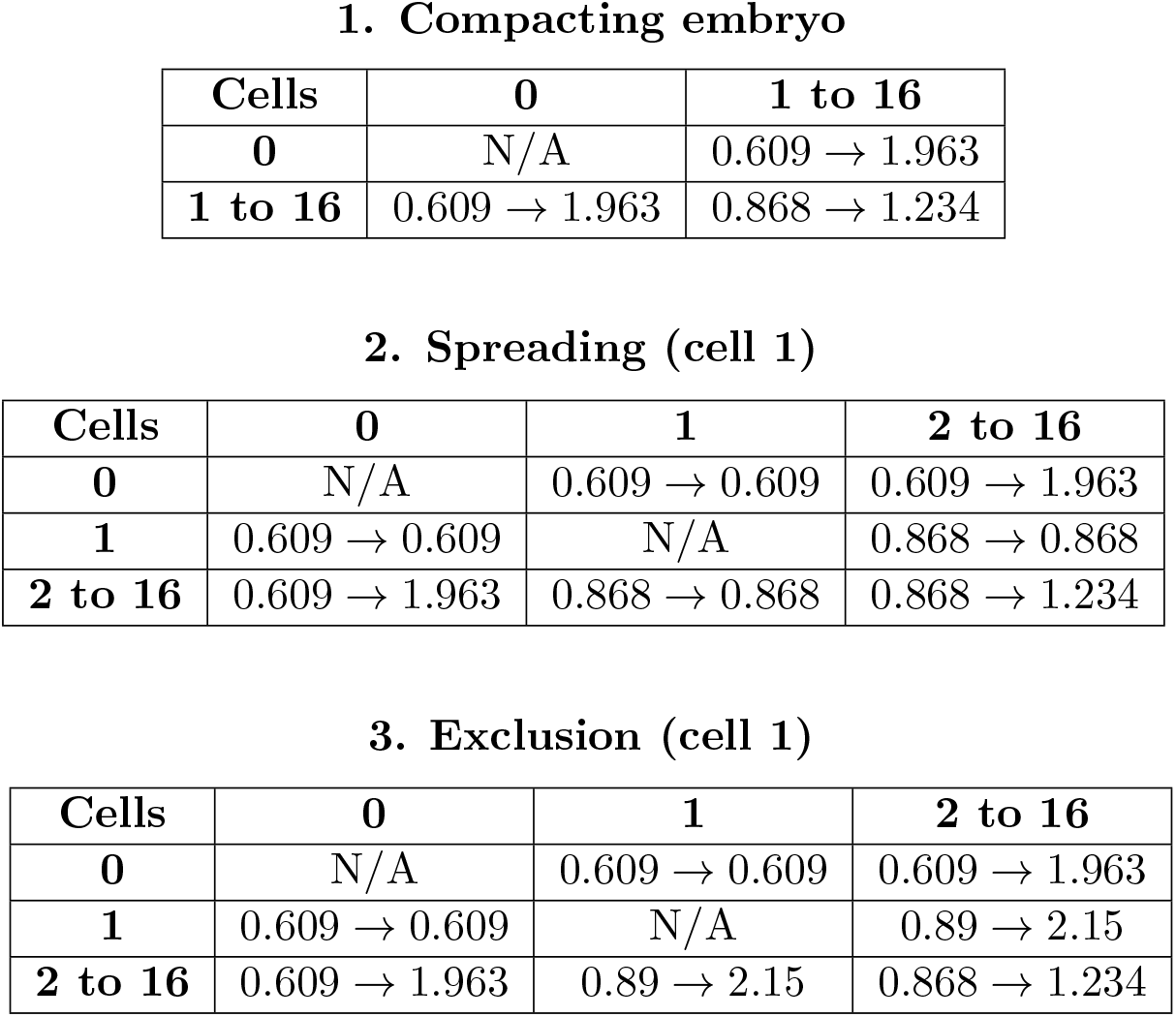
Tensions values used for simulations. Units correspond to 1000 pN/*μ*m.

## References

1. Firmin, J. & Maître, J.-L. Morphogenesis of the human preimplantation embryo: bringing mechanics to the clinics. Seminars in Cell & Developmental Biology S1084952121001920 (2021) doi:10.1016/j.semcdb.2021.07.005.

2. Shahbazi, M. N. Mechanisms of human embryo development: from cell fate to tissue shape and back. Development 147, dev190629 (2020).

3. Coticchio, G., Lagalla, C., Sturmey, R., Pennetta, F. & Borini, A. The enigmatic morula: mechanisms of development, cell fate determination, self-correction and implications for ART. Human Reproduction Update 25, 422–438 (2019).

4. Lagalla, C. et al. Embryos with morphokinetic abnormalities may develop into euploid blastocysts. Reproductive BioMedicine Online 34, 137–146 (2017).

5. Collinet, C. & Lecuit, T. Programmed and selforganized flow of information during morphogenesis. Nat Rev Mol Cell Biol (2021) doi:10.1038/s41580-020-00318-6.

6. Heisenberg, C.-P. & Bellaïche, Y. Forces in tissue morphogenesis and patterning. Cell 153, 948–962 (2013).

7. Haniffa, M. et al. A roadmap for the Human Developmental Cell Atlas. Nature 597, 196–205 (2021).

8. Rossant, J. & Tam, P. P. L. Opportunities and challenges with stem cell-based embryo models. Stem Cell Reports 16, 1031–1038 (2021).

9. Özgüç, Ö. & Maître, J.-L. Multiscale morphogenesis of the mouse blastocyst by actomyosin contractility. Current Opinion in Cell Biology 66, 123–129 (2020).

10. Fogarty, N. M. E. et al. Genome editing reveals a role for OCT4 in human embryogenesis. Nature 550, 67–73 (2017).

11. Gerri, C. et al. Initiation of a conserved trophectoderm program in human, cow and mouse embryos. Nature 587, 443–447 (2020).

12. Okamoto, I. et al. Eutherian mammals use diverse strategies to initiate X-chromosome inactivation during development. Nature 1–7 (2011) doi:10.1038/nature09872.

13. Petropoulos, S. et al. Single-Cell RNA-Seq Reveals Lineage and X Chromosome Dynamics in Human Preimplantation Embryos. Cell 165, 1012–1026 (2016).

14. Iwata, K. et al. Analysis of compaction initiation in human embryos by using time-lapse cinematography. J Assist Reprod Genet 31, 421–426 (2014).

15. Coticchio, G. et al. Perturbations of morphogenesis at the compaction stage affect blastocyst implantation and live birth rates. Human Reproduction 36, 918–928 (2021).

16. Rienzi, L. et al. Time of morulation and trophectoderm quality are predictors of a live birth after euploid blastocyst transfer: a multicenter study. Fertility and Sterility 112, 1080–1093.e1 (2019).

17. Skiadas, C., Jackson, K. & Racowsky, C. Early compaction on day 3 may be associated with increased implantation potential. Fertility and Sterility 86, 1386–1391 (2006).

18. Turlier, H. & Maître, J.-L. Mechanics of tissue compaction. Seminars in Cell & Developmental Biology 47–48, 110–117 (2015).

19. Goel, N. S., Doggenweiler, C. F. & Thompson, R. L. Simulation of cellular compaction and internalization in mammalian embryo development as driven by minimization of surface energy. Bulletin of mathematical biology 48, 167–187 (1986).

20. Maître, J.-L., Niwayama, R., Turlier, H., Nédélec, F. & Hiiragi, T. Pulsatile cell-autonomous contractility drives compaction in the mouse embryo. Nature cell biology 17, 849–855 (2015).

21. Schliffka, M. F. et al. Multiscale analysis of single and double maternal-zygotic Myh9 and Myh10 mutants during mouse preimplantation development. eLife 10, e68536 (2021).

22. Maître, J.-L. & Heisenberg, C.-P. Three Functions of Cadherins in Cell Adhesion. Current Biology 23, R626–R633 (2013).

23. Zakharova, E. E., Zaletova, V. V. & Krivokharchenko, A. S. Biopsy of Human Morula-Stage Embryos: Outcome of 215 IVF/ICSI Cycles with PGS. PLoS ONE 9, e106433 (2014).

24. Maitre, J.-L. et al. Adhesion Functions in Cell Sorting by Mechanically Coupling the Cortices of Adhering Cells. Science 338, 253–256 (2012).

25. Guck, J. Some thoughts on the future of cell mechanics. Biophys Rev 11, 667–670 (2019).

26. Budczies, J. et al. Cutoff Finder: A Comprehensive and Straightforward Web Application Enabling Rapid Biomarker Cutoff Optimization. PLoS ONE 7, e51862 (2012).

27. Coorens, T. H. H. et al. Inherent mosaicism and extensive mutation of human placentas. Nature (2021) doi:10.1038/s41586-021-03345-1.

28. Maître, J.-L. et al. Asymmetric division of contractile domains couples cell positioning and fate specification. Nature 536, 344–348 (2016).

29. Matamoro-Vidal, A. & Levayer, R. Multiple Influences of Mechanical Forces on Cell Competition. Current Biology 29, R762–R774 (2019).

30. True, J. R. & Haag, E. S. Developmental system drift and flexibility in evolutionary trajectories. Evol Dev 3, 109–119 (2001).

31. Lenne, P.-F. et al. Roadmap for the multiscale coupling of biochemical and mechanical signals during development. Phys. Biol. 18, 041501 (2021).

32. Clark, A. T. et al. Human embryo research, stem cell-derived embryo models and in vitro gametogenesis: Considerations leading to the revised ISSCR guidelines. Stem Cell Reports 16, 1416–1424 (2021).

33. Tsunoda, Y., Yasui, T., Nakamura, K., Uchida, T. & Sugie, T. Effect of cutting the zona pellucida on the pronuclear transplantation in the mouse. J Exp Zool 240, 119–125 (1986).

34. Guevorkian, K. Micropipette aspiration: A unique tool for exploring cell and tissue mechanics in vivo. Methods in cell biology 139, 187–201 (2017).

35. Schindelin, J. et al. Fiji: an open-source platform for biological-image analysis. Nat Methods 9, 676–682 (2012).

36. Alpha Scientists in Reproductive Medicine and ESHRE Special Interest Group of Embryology et al. The Istanbul consensus workshop on embryo assessment: proceedings of an expert meeting. Human Reproduction 26, 1270–1283 (2011).

## References

1. Clark, A. G., Wartlick, O., Salbreux, G. & Paluch, E. K. Stresses at the Cell Surface during Animal Cell Review Morphogenesis. Curr. Biol. 24, R484–R494 (2014).

2. Maître, J. L., Niwayama, R., Turlier, H., Nédélec, F., & Hiiragi, T. Pulsatile cell-autonomous contractility drives compaction in the mouse embryo. Nat. Cell Biol. 17, 849–855 (2015).

3. Maître, J. L., Turlier, H., Illukkumbura, R., Eismann, B., Niwayama, R., Nédélec, F., & Hiiragi, T. Asymmetric division of contractile domains couples cell positioning and fate specification. Nature 536, 344–348 (2016).

4. Turlier, H., Audoly, B., Prost, J. & Joanny, J.-F. Furrow constriction in animal cell cytokinesis. Biophys. J. 106, 114–123 (2014).

5. Khalilgharibi, N., Fouchard, J., Asadipour, N., Barrientos, R., Duda, M., Bonfanti, A., … & Charras, G. Stress relaxation in epithelial monolayers is controlled by the actomyosin cortex. Nat. Phys 15, 839–847 (2019).

6. Turlier, H., & Maître, J-L. Mechanics of tissue compaction. Seminars in cell & developmental biology 47, 110–117 (2015).

7. Nocedal, J., & Wright, S. Numerical optimization. (Springer Science & Business Media, 2006).

8. Hestenes, M. R., & Stiefel, E. Methods of conjugate gradients for solving linear systems. Washington, DC: NBS 59(1952).

9. Da, F., Batty, C., & Grinspun, E. Multimaterial mesh-based surface tracking. ACM Trans. Graph. 33, 112:1-112:11 (2014).

